# Dynlt3, a novel fundamental regulator of melanosome movement, distribution, maturation and transfer

**DOI:** 10.1101/2021.01.13.426541

**Authors:** Zackie Aktary, Alejandro Conde-Perez, Florian Rambow, Mathilde Di Marco, François Amblard, Ilse Hurbain, Graça Raposo, Cédric Delevoye, Sylvie Coscoy, Lionel Larue

**Author notes:** Address correspondence to Lionel LARUE, - - Institut Curie – Bat 110 - 91405, Orsay Cedex, France.

## Abstract

Skin pigmentation is dependent on cellular processes including melanosome biogenesis, transport, maturation and transfer to keratinocytes. However, how the cells finely control these processes in space and time to ensure proper pigmentation remains unclear. Here, we show that a component of the cytoplasmic dynein complex, Dynlt3, is required for efficient melanosome transport, maturation and transfer. In melanocytes with decreased levels of Dynlt3, pigmented melanosomes undergo a more directional (convective) motion leading to their peripheral location in the cell, and are not fully matured. Stage IV melanosomes are more acidic, but still heavily pigmented, resulting in a less efficient melanosome transfer. Finally, the level of Dynlt3 is dependent on β-catenin activity, revealing a novel function of the Wnt/β-catenin signalling pathway during melanocyte and skin pigmentation, by coupling the transport, position and maturation of melanosomes required for their transfer.

## INTRODUCTION

Melanocytes are neural crest derived cells that are primarily found in the epidermis and hair follicles, but are also found in the inner ear, eye, heart and meninges ^1^. These cells contain specialized organelles, known as melanosomes, which are responsible for pigmentation. Melanosomes belong to the lysosome related organelle (LRO) family and exist in four different stages ^2^. Only stage III and IV melanosomes are pigmented. Stage I melanosomes, the most primitive melanosomes, are membrane bound early endosomal structures that initiate the intraluminal Pmel fibrillation. The Pmel fibrils are fully formed in stage II melanosomes, following which melanin synthesis begins. In stage III, different melanogenesis enzymes (e.g. Tyrp1, Dct) are transported and inserted into the melanosome limiting membrane to produce melanin, which deposits on the pre-existing Pmel fibrils. Melanin accumulates in stage IV melanosomes that will become heavily pigmented. Melanosomes/melanin are then transferred to keratinocytes through an exocytosis/phagocytosis process to fulfill their primary role in protecting the skin from UV irradiation ^3^. Prior to melanosome exocytosis and melanin transfer, melanosomes must acquire a secretory signature through maturation steps that are still poorly understood. Maturation is a complex process that includes the melanosome lumen deacidification or neutralization, the acquisition of specific endosomal components by the melanosomes, and the concomitant removal of melanosome-associated membrane proteins (Vamp7 and most likely Tyrp1) through the generation and release of melanosomal membrane tubules ^4-6^.

The intracellular trafficking of melanosomes, similar to other LROs, involves the action of different families of motor proteins. Along the microtubules, melanosomes are transported by both kinesin and dynein motors ^7^. Most kinesins move their respective cargoes from the nucleus towards the periphery, whereas dynein motors move their cargoes in the opposite direction ^8^. At the cell periphery, melanosomes are transferred to the actin microfilaments, where the complex consisting of Myosin Va, Rab27a and Melanophilin move the melanosomes centrifugally to be transferred. While mutations in these three genes lead to pigmentation defects in humans and in mice, to date, no mutation in dynein and kinesin genes has been described in pigmentation.

In humans and mice, abnormal skin pigmentation results from a reduction/lack of melanocytes, a defect in melanin synthesis, and/or disrupted melanosome transport/maturation/transfer ^9^. We have previously shown that alterations of β-catenin activity (gain [bcat*] and loss of [Δex2-6]) result in the reduction of melanocyte proliferation and a coat color phenotype in mice ^10,11^.

In the current study, using melanocyte cell lines established from the skin of wild-type and bcat* C57BL/6J mice in culture, we observed a marked diminution of the number of pigmented melanosomes in the perinuclear area of bcat* melanocytes. We identified Dynlt3, a member of the cytoplasmic dynein complex as a downregulated component in bcat* melanocytes compared to wild-type, and as a regulator of melanosome movement, distribution, maturation and transfer. Our data demonstrate, for the first time, the fundamental role of Dynlt3 in coat and skin coloration.

## RESULTS

### Melanocytes expressing stable β-catenin display peripheral melanosome distribution

We established melanocyte cell lines in culture from the skin of wild-type (9v) and bcat* (10d) C57BL/6J pups. As expected, WT pigmented melanosomes (mainly stages III and IV) were distributed uniformly throughout the cell (Fig. 1a,b, left).

**Figure 1.**
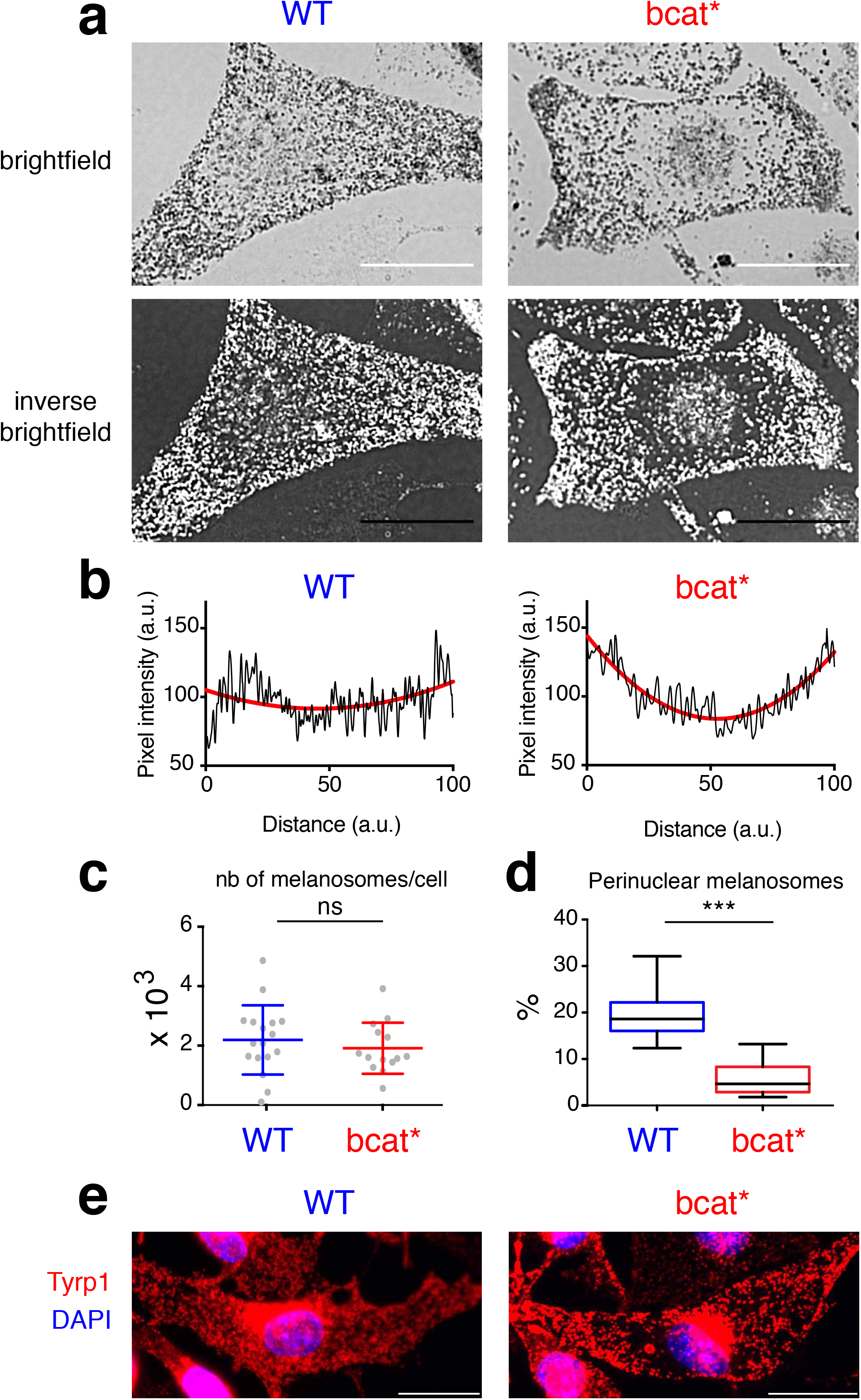
Melanosomes are distributed at the periphery of melanocytes expressing activated β-catenin. a) Brightfield (top) and inverse brightfield (bottom) images of WT (9v) and bcat* (10d) melanocytes. Bar, 20 μm. b) Quantitation of pixel intensity (in black) across the length of the WT and bcat* melanocytes shown in Panel A. Trendlines are shown in red. c) Quantification of the total number of melanosomes in WT and bcat* melanocytes. Results represent the mean ± SD of pooled data from three independent experiments with a total of nineteen WT and fourteen bcat* cells. ns signifies no statistical significance (p > 0.05) as determined by the two-sided Mann-Whitney test. d) Quantification of the number of perinuclear melanosomes (in a 15 µm diameter circle centered around the nucleus) in WT and bcat* melanocytes. Results represent the mean ± SD of pooled data from three independent experiments with a total of nineteen WT and fourteen bcat* cells. Statistical significance was determined by the two-sided Mann-Whitney test and *** signifies p < 0.001. e) Immunofluorescence analysis of WT and bcat* melanocytes. Cells were processed for immunofluorescence and stained with anti-Tyrp1 (red) antibodies and the nuclei were stained with DAPI (blue). Note that Tyrp1 staining was also observed in compartments adjacent to the nucleus, which is most likely the protein that is being synthesized and processed in the Golgi/ER. Bar, 20 μm.

However, melanosome localization was markedly different in bcat* melanocytes, which displayed a notable absence of pigmented melanosomes in the perinuclear area (Fig. 1a,b, right), while the total number of melanosomes in WT and bcat* cells was similar (Fig. 1c). The quantification of the number of perinuclear melanosomes in a 15 μm area around the nucleus revealed that ≈ 20% of the pigmented melanosomes were perinuclear in WT cells, compared to only 5% in bcat* cells (Fig. 1d). Immunofluorescence microscopy using anti-Tyrp1 antibodies-mainly labeling stages III and IV melanosomes ^12^- further confirmed the reduction of pigmented melanosomes in the perinuclear area of bcat* cells as compared to WT melanocytes (Fig. 1e).

To confirm that the observed reduction of perinuclear melanosomes in this mutant cell line was not limited to the specific bcat* construct, we expressed another active β-catenin mutant, the βcat-Δex3-GFP mutant, in WT melanocytes ^13^. This mutant construct encodes for a β-catenin protein that lacks the serine and threonine residues that allow for its proteasomal degradation. Similar to the bcat* cells, expression of βcat-Δex3-GFP in WT melanocytes (WT-βcatGFP, fluorescent cells) resulted in a reduced number of perinuclear melanosomes compared to a GFP control (WT-GFP, fluorescent cells) (Supplementary Fig. 1).

Pigmented melanosomes consist of stage III and IV melanosomes. Ultrastructurally, stage III melanosomes contain dark and thick intraluminal melanin-positive fibrils, while stage IV melanosomes are fully filled with pigment. Melanosome maturation was evaluated in WT and bcat* cells by transmission electron microscopy (TEM) assay. Intriguingly, pigmented bcat* melanosomes were significantly larger than in WT melanosomes (≈370 nm *vs*. ≈300 nm) (Supplementary Fig. 2).

### β-catenin affects Dynlt3 expression

The notable peripheral melanosome distribution in bcat* cells very likely reflected a disruption in their proper intracellular trafficking, due to the exogenous expression of β-catenin. To examine this possibility, we looked at the transcriptome of WT and bcat* melanocytes and specifically focused on the expression of 67 genes known to be involved in Lysosomal Related Organelle (LRO) biogenesis, transport and maturation (Fig. 2a and Table 1). From these analyses, we identified Dynlt3, a member of the cytoplasmic dynein complex, as a gene whose expression was decreased two-fold in bcat* cells compared to WT cells.

**Figure 2.**
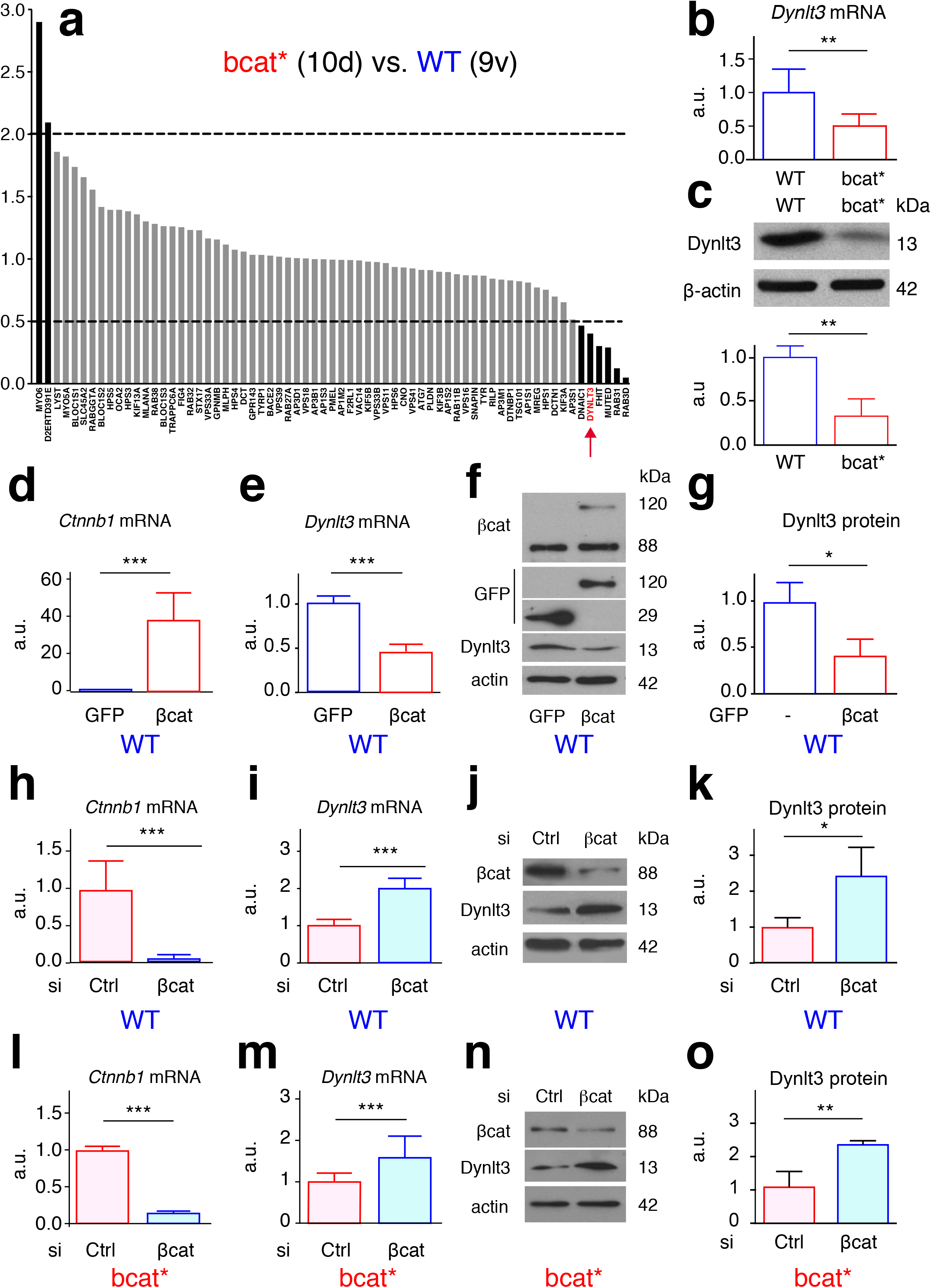
Dynlt3 is downregulated after β-catenin expression. a) Expression profile of 67 genes associated with trafficking of melanosomes and other lysosome related organelles (LRO). Data was obtained from Affymetrix mRNA transcriptomic analyses and is presented as the ratio of expression of bcat* (10d) cells to WT (9v) cells. The raw data set is available upon request. b) RT-qPCR analysis of Dynlt3 mRNA levels in WT and bcat* melanocytes. c) Western blot analysis of Dynlt3 protein levels in WT and bcat* melanocytes. Quantification of the data is presented below from four independent experiments. d) d,e) RT-qPCR analysis of β-catenin (D) and Dynlt3 (E) expression in WT melanocytes following transfection with the βcat-Δex3-GFP (βcat) expression vector or a GFP control. f) Western blot of βcat and Dynlt3 levels in WT melanocytes following transfection with the βcat-Δex3-GFP (βcat) expression vector or a GFP control. g) Quantification of the Dynlt3 western blot data presented in Panel F. h,i) RT-qPCR analysis of β-catenin (H) and Dynlt3 (I) expression in WT melanocytes following transfection with either a control siRNA or siRNA targeting β-catenin. j) Western blot of βcat and Dynlt3 levels in WT melanocytes following transfection with either a control siRNA or siRNA targeting β-catenin. k) Quantification of the Dynlt3 western blot data presented in Panel J. l,m) RT-qPCR analysis of β-catenin (L) and Dynlt3 (M) expression in bcat* melanocytes following transfection with either a control siRNA or siRNA targeting β-catenin. n) Western blot of βcat and Dynlt3 levels in bcat* melanocytes following transfection with either a control siRNA or siRNA targeting β-catenin. o) Quantification of the Dynlt3 western blot data presented in Panel N. For western blot analysis, all statistical significance was determined using unpaired two-sided T-tests. *: p < 0.05; **: p < 0.01; ***: p < 0.001.

Transcriptomic data were confirmed by RT-qPCR and western blot analyses and showed that both Dynlt3 mRNA and protein levels were decreased in bcat* cells, relative to WT cells (Fig. 2b,c). These results were corroborated using independent melanocyte cell lines established from independent WT and bcat* pups (Supplementary Fig. 3a). The regulation of Dynlt3 levels by β-catenin was then validated using different approaches. We first overexpressed βcat-Δex3-GFP in WT cells, following which Dynlt3 mRNA and protein levels were both significantly decreased (Fig. 2d-g). Subsequently, we knocked down β-catenin in WT cells and observed an increase in both Dynlt3 mRNA and protein (Fig. 2h-k). Finally, the knockdown of β-catenin in bcat* melanocytes resulted in a similar increase in Dynlt3 mRNA and protein levels (Fig. 2l-o), thereby clearly demonstrating that β-catenin expression appears to have an inhibitory effect on the levels of Dynlt3.

### Knockdown of *Dynlt3* in WT melanocytes phenocopies melanosome distribution in bcat* melanocytes

Given that Dynlt3 is a member of the Tctex-type 3 family of Dynein motor light chain implicated in retrograde transport ^14,15^ and that its expression was decreased in bcat* cells, we wondered whether Dynlt3 was a player involved in melanosome distribution. To this end, we reduced the levels of *Dynlt3* in WT melanocytes. Similar to bcat* cells, knockdown of *Dynlt3* phenocopied the exclusion of pigmented melanosomes from the perinuclear area (Fig. 3a-c). The number of perinuclear melanosomes was decreased two-fold in WT-siDynlt3 cells as compared to control (Fig. 3d), while the total number of melanosomes was unchanged (Fig. 3e). As previously, WT-siDynlt3 cells stained with an anti-Tyrp1 antibody revealed a notable absence of staining in the perinuclear area, which was not the case in WT-siCtrl melanocytes (Fig. 3f). To confirm these results, we overexpressed Dynlt3 in bcat* melanocytes, which resulted in the redistribution of the melanosomes in the perinuclear area (Fig. 4).

**Figure 3.**
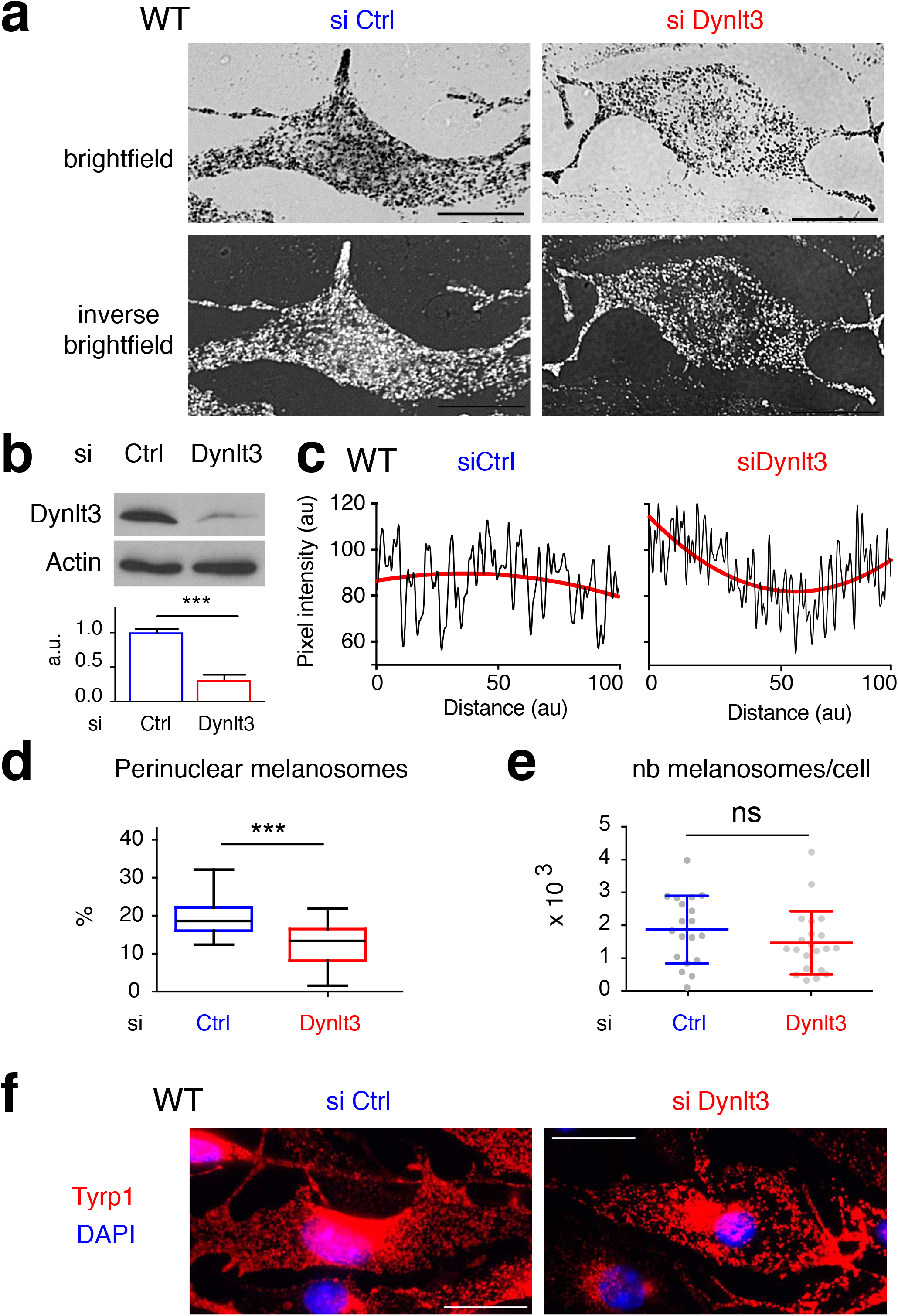
Melanosomes are distributed primarily at the periphery of melanocytes when Dynlt3 levels are decreased. a) Brightfield (top) and inverse brightfield (bottom) images of WT (9v) cells transfected with either siControl (left) or siDynlt3 (right). Bar, 20 μm. Ctrl means control. b) Western blot analysis of Dynlt3 levels in WT cells transfected with siControl (Ctrl) or siDynlt3. Relative quantification of the western blot analysis was performed from three independent experiments. Statistical significance was determined using unpaired two-sided T-tests. *** p<0.002. c) Quantitation of pixel intensity (in black) across the length of the WT cells transfected with siControl or siDynlt3 shown in Panel A. Trendlines are shown in red. d) Quantification of the number of perinuclear melanosomes (in a 15 µm diameter circle centered around the nucleus) in WT melanocytes transfected with siCtrl or siDynlt3. Results represent the mean ± SD of pooled data from three independent experiments with a total of nineteen WT-siCtrl and twenty-one WT-siDynlt3 cells. Statistical significance was determined by the two-sided Mann-Whitney test, ***: p < 0.001. e) Quantification of the total number of melanosomes in WT melanocytes transfected with siCtrl or siDynlt3. Results represent the mean ± SD of pooled data from three independent experiments with a total of nineteen WT-siCtrl and twenty-one WT-siDynlt3 cells. Statistical significance was determined by the two-sided Mann-Whitney test, ns: not significant. f) Immunofluorescence analysis of WT melanocytes transfected with siCtrl or or siDynlt3. Cells were processed for immunofluorescence 48 hours after transfection and stained with anti-Tyrp1 (red) antibodies. Nuclei were stained with DAPI (blue). Note that Tyrp1 staining was also observed in compartments adjacent to the nucleus, which is most likely the protein that is being synthesized and processed in the Golgi/ER. Scale bar, 20 μm.

**Figure 4.**
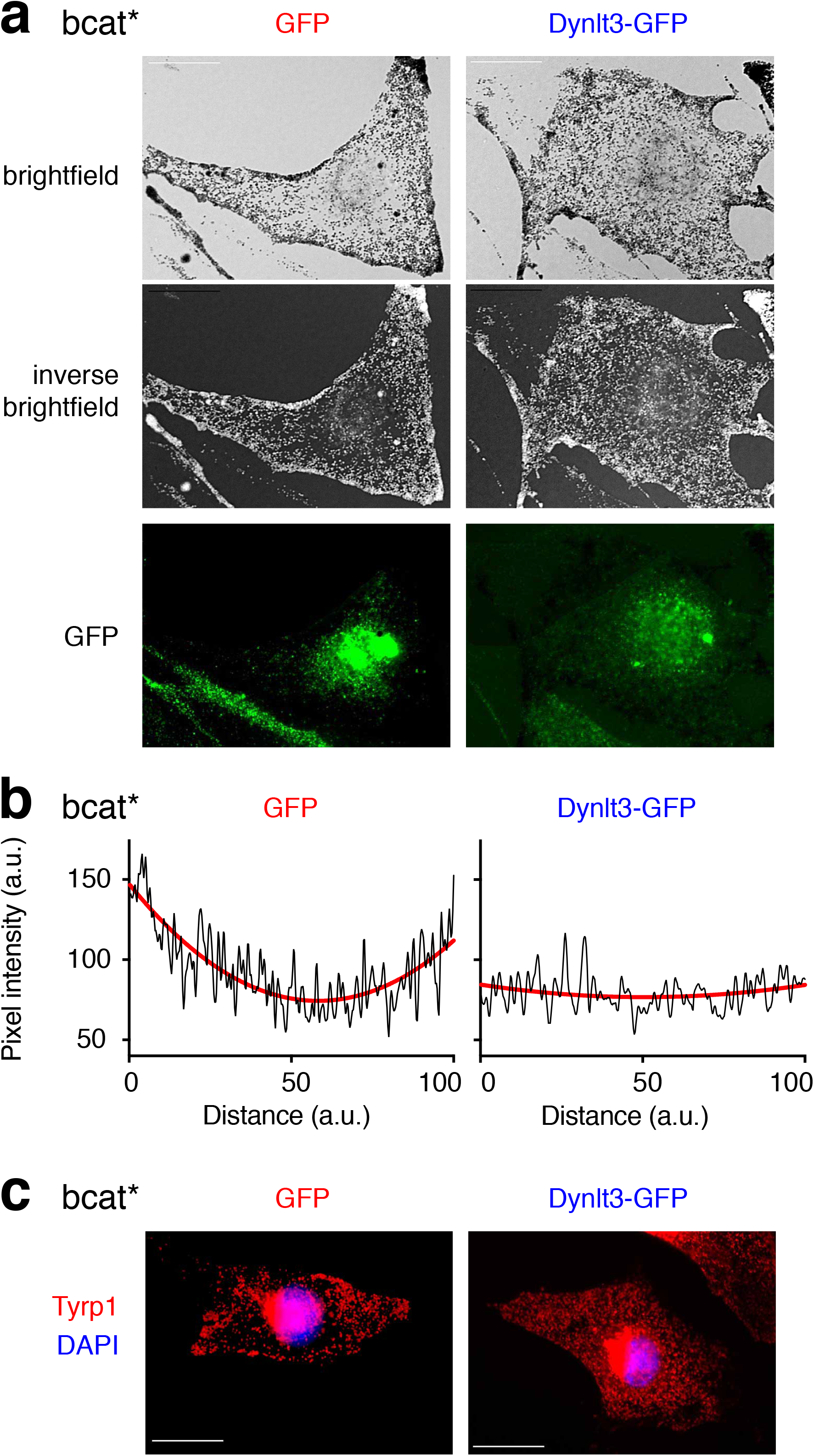
Overexpression of Dynlt3 redistributes melanosomes in bcat* melanocytes. a) Brightfield (top), inverse brightfield (middle) and GFP (bottom) images of bcat* (10d) melanocytes transiently expressing GFP or GFP-tagged Dynlt3. Bar, 20 μm. b) Quantitation of pixel intensity (in black) across the length of the bcat*-GFP and bcat*-Dynlt3GFP melanocytes shown in Panel A. Trendlines are shown in red. c) Immunofluorescence analysis of bcat* melanocytes transiently expressing GFP or GFP-tagged Dynlt3. Cells were processed for immunofluorescence and stained with anti-Tyrp1 (red) antibodies and the nuclei were stained with DAPI (blue). Bar, 20 μm.

### Decreased Dynlt3 amounts increased melanosome motility

We next examined potential alterations in melanosome motility using time-lapse video microscopy to track pigmented melanosome movement. Direct observation of melanosome movements revealed that generally motion is composed of variable and intermittent movement, with high level of pauses followed by short bursts of movements, as previously reported ^16,17^.

The movement of melanosomes in bcat* cells was enhanced when compared with WT melanosomes. On average, the total distance that bcat* melanosomes traveled was longer than WT melanosomes (24.0 μm *vs.* 11.8 μm, Fig. 5a,b). Similarly, βcat-⊗ex3-GFP WT cells showed increased total distance travelled by melanosomes relative to WT cells expressing GFP alone (21.4 μm *vs.* 11.1 μm, Fig. 5a,b). Additionally, following transfection with GFP or βcat-⊗ex3-GFP, we monitored the trajectories of melanosomes in GFP negative cells (i.e. that were unsuccessfully transfected). The total distance that the melanosomes travelled on average in these two non-GFP cells (GFP or βcat-⊗ex3-GFP) was similar to WT cells (11.8 μm vs. 11.0 μm vs. 11.3 μm, respectively, Fig. 5a,b). Taken together, β-catenin exogenous expression increased movement of melanosomes.

**Figure 5.**
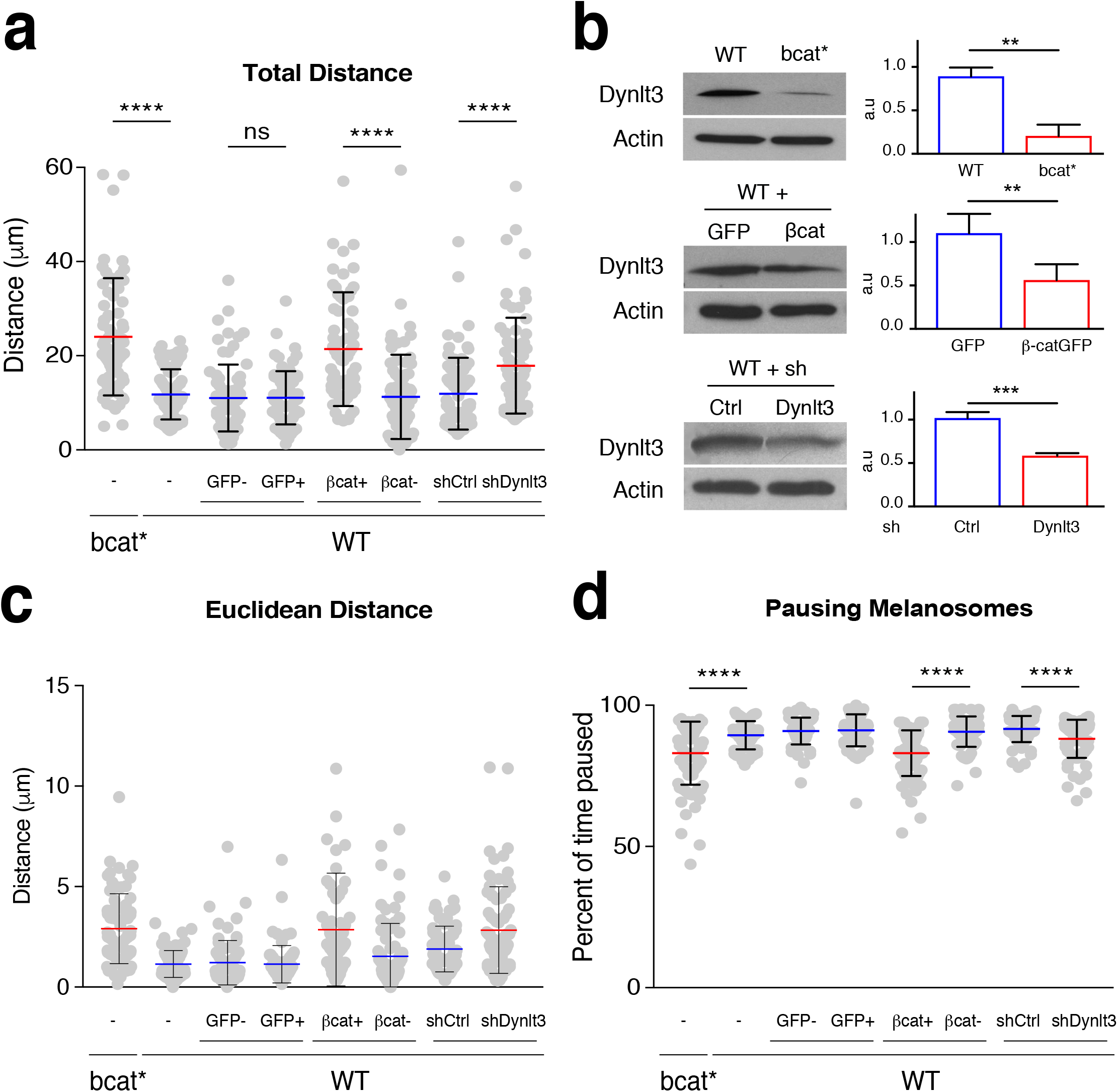
Melanosome movement is increased in cells producing active β-catenin or with diminished levels of Dynlt3. Melanosome movement (corresponding mainly to stage IV melanosome) was assessed by brightfield video microscopy over a period of 5 minutes and melanosome trajectories were followed using *ImageJ* software. For each cell line, 75 melanosomes were followed from a minimum of 5 independent cells. WT (9v) cell lines were transfected with control GFP or β-catenin-GFP expression vectors, respectively. Analyses were performed on both transfected (i.e. GFP-positive) and non-transfected (i.e. GFP-negative) cells. In addition, WT cells were transfected with either control or Dynlt3 shRNA vectors and analyses were performed on the resulting red cells. Each dot represents one melanosome. (a) The total distance indicates the sum of all of the individual tracks for each melanosome, (c) The Euclidean distance refers to the distance between the start and end position of each melanosome. (d) The percentage of time that each melanosome spent stationary/paused refers to the proportion of the number of tracks where no movement was measured compared to the total number of tracks. (b) The level of Dynlt3 in these different cell lines was evaluated after western blot analysis. Relative quantification of the western blot analysis was performed from three to four independent experiments. In panel c, one outlier was removed from WT+βcat+, its value was 20 μm. For western blot analysis, all statistical significance was determined using unpaired two-sided T-tests. **:p <0.01; *** p<0.001. For the other analyses, statistical significance was measured using two-sided Mann-Whitney test. **** p < 0.0001.

Similarly, downregulation of Dynlt3 in WT melanocytes stably expressing shRNA targeting this gene resulted in a significant decrease of Dynlt3 and an increase in the total distance melanosomes travelled compared to shCtrl cells (17.9 μm *vs.* 11.9 μm, Fig. 5a,b), accompanied by an increased average distance and average velocity of melanosomes (Supplementary Fig. 4). Consistently, Euclidean distance travelled by the melanosomes was increased after overexpression of β-catenin or downregulation of Dynlt3 (Fig. 5b,c).

Finally, the percentage of pausing melanosomes was determined by considering the number of frames where no movement was recorded. Tracked melanosomes in bcat*, WT-βcat-⊗ex3 or WT-shDynlt3 cells paused less frequently than controls (Fig. 5b,d), showing together that Dynlt3, through bcat expression, contributes to at least the intracellular dynamic of pigmented melanosomes.

### Mathematical and statistical analyses reveal a convective melanosome movement after reduction of Dynlt3 levels

Displacement of an object can be generated by a superposition of various elementary processes: Brownian movement (diffusion) as a basis process, with the potential addition of velocity-driven movement (convection or directed movement), elastic tethering (flexible attachment to a fixed point) and/or confinement. “Not convective” conditions refer here to populations of trajectories that could be generated without addition of a velocity-driven movement (Fig. 6a,b). In order to identify the elementary physical processes involved in melanosome movement, we compared experimental melanosome trajectories (x, y, t) with simulated trajectories integrating these different types of movements ^18^. This comparison is based on eight spatio-temporal descriptors (see supplementary information). Here, we focus on a descriptor referring to the classical analysis of the Mean-square displacement (MSD or <r^2^>) during a time interval (t).

**Figure 6.**
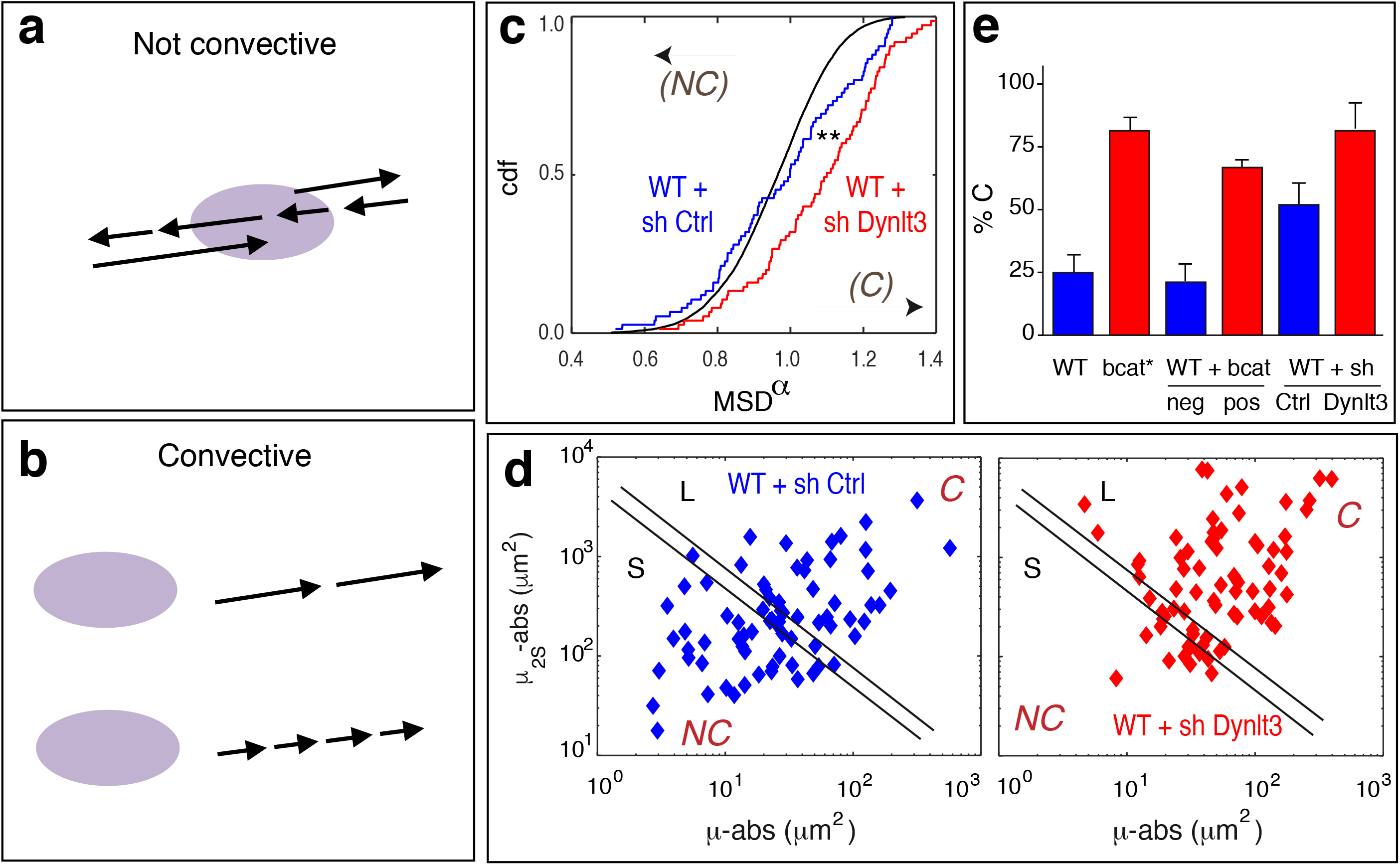
Increased convective movement of melanosomes in cells with increased levels of β-catenin or decreased Dynlt3. a,b) Schematic representation of not convective (a) and convective (b) movement. The purple ellipse represents the cargo. The arrows represent the direction and the orientation of the movement, and the length of the arrows are directly proportional to the magnitude of the movement. c) α MSD distributions for WT-shCtrl (blue) and WT-shDynlt3 (red) populations of trajectories, compared to the distribution for simulated Brownian (purely diffusive) random walks (nwalks=10 000, with the same nsteps (601) as for experimental trajectories). Theoretical shifts compared to Brownian process when adding convection, noise, tethering or confinement are indicated with arrows. **, statistically significant difference between WT-shDynt3 and WT-shCtrl, p<0.005. Statistical analyses were done using a Wilcoxon test. d) µ_2S_,µ_2L_ plots reflecting the extent of WT-shCtrl (b) and WT-shDynlt3 (c) trajectories. Each point corresponds to one trajectory. Lines correspond to frontiers between non-convective (NC) and convective (C) trajectories: see SI for criteria and validation. e) Number of non-convective (NC) trajectories for melanosomes in WT, bcat*, WT-GFP, WT-βcat-Δex3-GFP, WT-shCtrl and WT-shDynlt3 cells, out of 75 trajectories for each population of trajectories. The same frontiers were used for all populations (see SI). Error bars correspond to trajectories between the two frontiers (see Fig. 5d, and supplementary Fig. 5S).

In the case of purely 2D diffusive motion, MSD(t) = 4Dt (D, diffusion coefficient), while for purely directed movement MSD(t) = V^2^t^2^ (V, velocity). Here, the MSD was fit with <r^2^> = *bt*^α^, and α, the MSD exponent, was taken as our main descriptor (Fig. 6c, α cumulative distribution function, cdf). α gives access to the convective (α shifts towards 2) or not convective (α centered about 1) characteristics of the movement.

Compared to a Brownian reference process (black), half of WT-shCtrl trajectories (blue) were shifted to the left and correspond to non-convective trajectories, while the other half were shifted to the right, indicative of convective movement (Fig. 6d). The vast majority of WT-shDynlt3 trajectories (red) were convective (Fig. 6d), with a statistically significant difference between shCtrl and shDynlt3 α distributions (p<0.005, Wilcoxon test). Similarly, there was a statistically significant difference between the α distributions of WT and bcat* cells and between WT-GFP and WT-βcat-Δex3-GFP cells (Supplementary Fig. 5), with the melanosomes in cells overexpressing β-catenin displaying more convective movement. Accordingly, the populations were subdivided in two subpopulations, small [S] and large [L] amplitudes, as assessed by the second moments of trajectories (Fig. 6d). The frontier was chosen such that S mainly corresponded to a homogeneous not convective (NC) population, and L to a convective (C) population (see supplementary information), with trajectories between the two lines defining an error bar. The number of non-convective trajectories (out of 75 trajectories) was estimated to be 56-62 for WT, compared to 14-18 for bcat*; 60-64 for WT-GFP compared to WT-βcat-Δex3-GFP (Supplementary Fig. 5); and 36-43 for WT-shCtrl compared to 14-22 for WT-shDynlt3 (Fig. 6e). This illustrates that the downregulation of Dynlt3 or the expression of active β-catenin both lead to a significant increase of convective trajectories.

### Overexpression of β-catenin or reduction of Dynlt3 diminishes melanosome maturation and transfer

Melanosome maturation was evaluated in cells expressing different amount of Dynlt3 by assessing melanosome acidity. Non-fully mature melanosomes are more acidic than mature melanosomes. In this context, the number of acidic and pigmented melanosomes was approximately two-fold higher in bcat* cells relative to WT cells (12.4 *vs.* 6.4 acidic melanosomes/field), as well as in WT-siDynlt3 *vs*. WT-siCtrl (13.0 *vs.* 7.3) (Fig. 7a).

**Figure 7.**
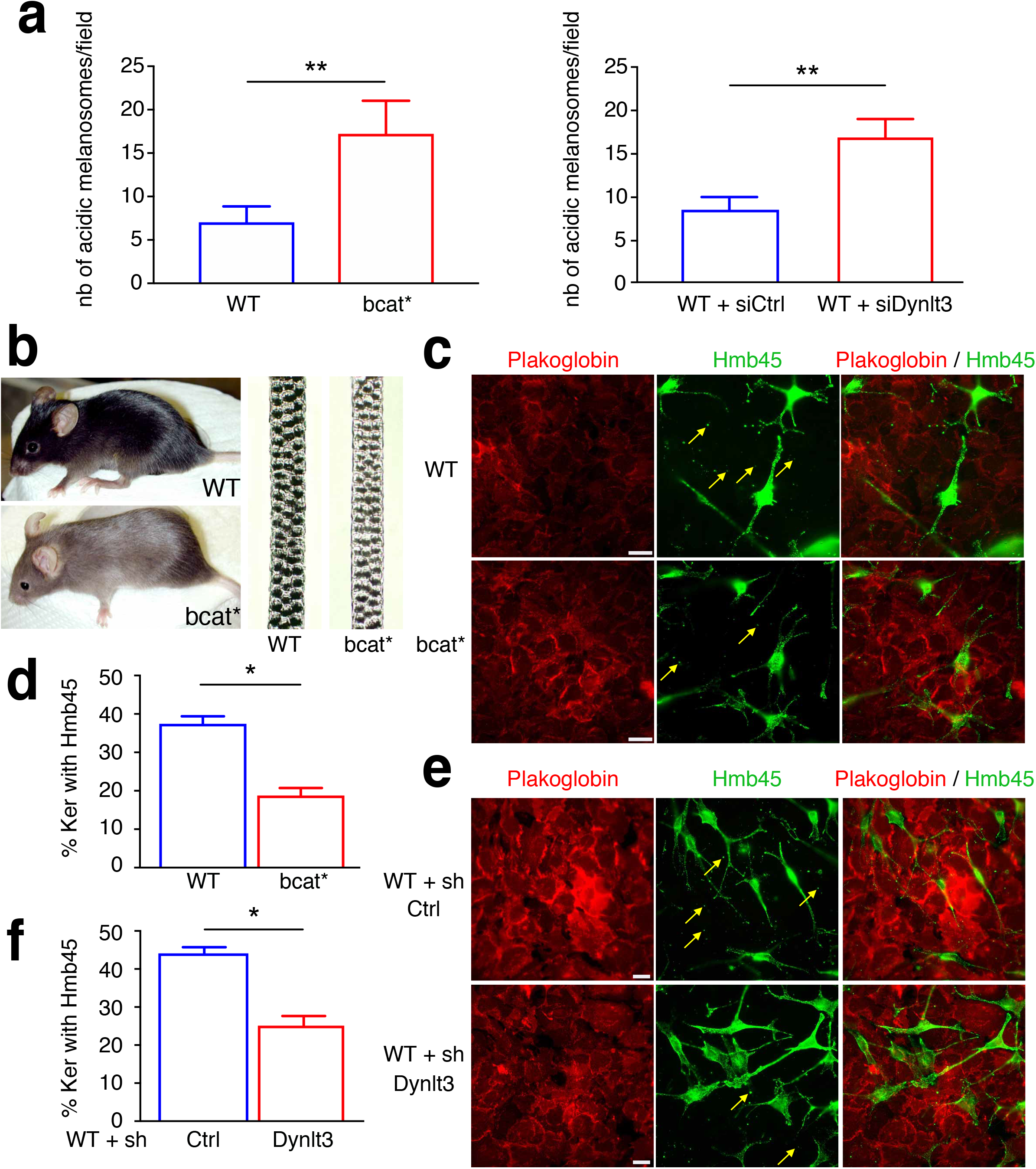
Melanosome maturation and transfer are altered in cells producing active β-catenin or with diminished level of Dynlt3. a) Quantitation of acridine orange positive melanosomes in WT, bcat*, WT + siCtrl and WT + si Dynlt3 cells. b) Photograph of wild-type and bcat* adult mice (left) and of their respective dorsal hairs (right). c) Immunofluorescence analysis of melanosome transfer between melanocytes WT (9v) or bcat* (10d) and Balb/c MK keratinocytes. Melanocytes and keratinocytes were co-cultured for 10 days, fixed and processed for immunofluorescence analyses using Plakoglobin and HMB45 antibodies. The yellow arrows indicate Pmel positive staining localized outside of melanocytes. d) Quantitation of the percentage of Pmel and Plakoglobin double positive cells is presented (right). Pmel positivity is assessed as being either positive or negative within each respective keratinocyte. Quantitation was done from a minimum of three independent co-cultures. e) Immunofluorescence analysis of melanosome transfer between melanocytes (WT + sh Ctrl or WT + shDynlt3) and Balb/c MK keratinocytes. Melanocytes and keratinocytes were co-cultured for 10 days, fixed and processed for immunofluorescence analyses using Plakoglobin and HMB45 antibodies. The yellow arrows indicate Pmel positive staining localized outside of melanocytes. f) Quantitation of the percentage of Pmel and Plakoglobin double positive cells is presented (right). Pmel positivity was assessed as being either positive or negative within each respective keratinocyte. Quantitation was done from a minimum of three independent co-cultures.

Melanocytes with increased β-catenin or decreased Dynlt3 levels harboured more pigmented and peripheral melanosomes, which would represent an ideal subpopulation of pigment granules to be transferred to keratinocytes. However, their change in pH might reflect a failure to acquire transfer capacity ^5^. We thus examined whether these cells would have a deficiency in melanosome transfer and consequently on coat color. Indeed, the hairs of bcat* C57BL/6J mice were lighter than WT C57BL/6J mice (Fig. 7b). Melanosome transfer was evaluated by co-culturing WT, bcat* WT-shCtrl, and WT-shDynlt3 melanocytes in the presence of Balb/c MK keratinocytes for 10 days prior to immunofluorescence analysis using plakoglobin (keratinocyte marker) and Hmb45 (a melanosome marker labeling the intraluminal melanin-positive fibrils) (Fig. 7c,e). We then counted the number of keratinocytes in which we could detect at least one Hmb45-positive structure; this number was significantly lower when Dynlt3 was lower than normal (Fig. 7d,f). In addition, we further confirmed that melanosome transfer was decreased in melanocytes with decreased Dynlt3 by FACS (Supplementary Fig. 6).

Taken together, these data suggest that in melanocytes with stable, activated β-catenin or in those with diminished expression of Dynlt3, both melanosome maturation and transfer are negatively affected.

## DISCUSSION

In this study, we showed that in melanocytes derived from transgenic mice expressing active β-catenin (bcat* cells), pigmented melanosomes were primarily absent from the perinuclear area and localized at the cell periphery. Notably, the expression of Dynlt3, a member of the cytoplasmic dynein complex, was downregulated in bcat* cells and knockdown of Dynlt3 in WT melanocytes phenocopied the peripheral localization of pigmented melanosomes and their absence from the perinuclear area. Melanosome movement was increased and melanosomes moved in a more convective (directed) manner in cells with diminished levels of Dynlt3. Decreased levels of Dynlt3 resulted in more acidic melanosomes that was associated with a reduction of melanosome transfer to keratinocytes. Altogether, these results show that Dynlt3 is a key player in regulating melanosome movement, maturation and transfer.

Dynlt3 is a component of the cytoplasmic dynein complex, a macromolecular protein complex that is essential for the transport of cargos from the cell periphery towards the nucleus ^14,15^. While the heavy chain (Dync1h1) of this complex has an ATPase activity and interacts with the microtubules, the light intermediate (Dync1li1 and Dync1li2) and the light chains, including Dynlt3, along with dynactin, a dynein-interacting cofactor and activator, are thought to play a role in cargo recognition.

Despite previous studies having shown that various virus capsid proteins (e.g. papilloma and herpes) interact with Dynlt3 and hijack the Cytoplasmic Dynein complex to reach the nucleus of infected cells ^19,20^, this work is the first to demonstrate functionally that Dynlt3 is involved in the intracellular trafficking of an organelle.

To date, transgenic mice with melanocyte-specific mutations in cytoplasmic dyneins have neither been identified nor produced, and therefore the role of Dynlt3 in mouse pigmentation remains unknown. However, transgenic mice containing heterozygous mutations in *Dync1h1*, which encodes the dynein heavy chain Dync1h1, have been produced. In general, these mice show defects in motor functions, with motor neuron degeneration and/or sensory neuropathy observed ^21-24^. Yet, no obvious coat color phenotypes were reported in these heterozygous mice. Importantly, homozygous Dync1h1 mice are not viable and die during embryonic development or shortly after birth. Therefore, whether a coat color phenotype would be observable in homozygous mice but would not apparent in heterozygous mice remains unknown. Conditional mouse mutants would be required to answer this point, using for instance the Tyr::Cre mice ^25^. While mutations in *DYNLT3* are not significantly detailed in human disease, reduction of DYNLT3 protein has been observed in late stage Parkinson’s disease that did not develop melanoma ^26^.

Microarray analyses of LRO trafficking genes demonstrated that among the components associated with cytoplasmic dynein, only Dynlt3 levels were altered in response to β-catenin expression, suggesting that only Dynlt3 is a target of β-catenin. Importantly, our results identify Dynlt3 as a novel β-catenin target, and one that whose expression is negatively regulated by β-catenin. However, we cannot exclude that knockdown of other dynein complex members may have a similar effect on melanosome localization. Importantly, in WT and bcat* cells, Dynlt1, the other member of the Tctex light chain family, was not expressed (Supplementary Fig. 2b), in agreement with previous studies showing that Dynlt1 and Dynlt3 are not expressed in the same tissue ^27,28^. Whether other cytoplasmic dynein subunits affect melanosome localization, and which ones in particular do so, are currently under investigation.

Video microscopy and manual tracking showed that melanosome movement is not directional (non-convective). Instead, the direction that melanosomes moved changed consistently, with melanosomes often going forwards and backwards repeatedly. Downregulation of Dynlt3 resulted in a more directed movement with changes in direction occurring less frequently. On microtubules, this back and forth movement of melanosomes can be thought of as competitions between the kinesin and dynein motors. While this random movement would be energetically costly and seemingly inefficient, this back and forth movement may occur to allow melanosomes to mature properly without passing onto the actin microfilaments and being transferred before they are fully mature.

To our knowledge, this work is the first indication that defects in a cytoplasmic dynein has an impact on melanosome transfer. That decreased Dynlt3 levels diminished melanosome transfer is somewhat perplexing since pigmented melanosomes localized primarily at the cell periphery, and were ideally positioned to be further transferred. However, in cells that expressed less Dynlt3, melanosomes were more acidic, as more pigmented melanosomes accumulated acridine orange. Since melanosomes de-acidify their lumen before being transferred to keratinocytes, and since increased acidity is an indication of immature melanosomes, our data suggests that Dynlt3 plays some role in the regulation of melanosome maturity, or at the very least, in the de-acidification of the melanosome lumen. Recently, it has been shown that the maturation of melanosomes before transfer involves the formation and release of membrane tubules containing proteins (e.g. Vamp7) that are needed to tune the melanosome homeostasis (including the pH and melanin content) and ultimately their secretory capacity ^5^. The separation of these tubules from the mature melanosome requires proteins involved in membrane scission (e.g. Myosin 6, Optineurin), whose downregulation results in increased melanosome size and acidity. One should note that Myosin 6 was upregulated in bcat* vs. WT melanocytes (Fig. 2a). Our hypothesis is that since the level of Dynlt3 is reduced, the cell tries to compensate for this reduction by increasing the amount of Myosin 6 to allow the separation of these tubules from the melanosomes. Therefore, it is conceivable that Dynlt3 plays a role in this process, possibly in the pulling, fission and/ or the retrograde transport of the melanosomal tubules, therefore contributing to melanosome maturation and secretion.

Taken together, our results have identified Dynlt3 as a novel and important player in melanosome biology (transport, maturation and transfer) and in coat coloration and pigmentation. Furthermore, our results suggest that cytoplasmic dynein proteins may have other roles in the biology of different organelles, in addition to their already established functions in organelle trafficking.

## MATERIALS AND METHODS

### Cell culture

Mouse primary melanocyte cell lines were established from the skin of wild-type (WT) and bcat* C57BL/6J transgenic mice as previously described ^10^. Melanocyte cell lines were grown in Ham’s F12 media (Gibco, 21765-029) supplemented with 10% FBS (Gibco, 10270-106), antibiotics (100 U/mL penicillin and 100 μg/mL streptomycin; Gibco, 15140), and 200nM TPA (Sigma, P1585). Balb/c MK mouse epidermal keratinocytes were grown in Joklik Modified MEM (Sigma, M8028) supplemented with 10% FBS, antibiotics (as above) and 5 ng/mL EGF (Sigma, E9644). All cell lines were maintained at 37°C in a humidified atmosphere containing 5% CO_2_.

For co-culture experiments, the melanocyte and keratinocyte cell lines were cultured in melanocyte growth media in the presence of 5 ng/mL EGF and 50 nM CaCl_2_. Co-culture experiments were performed with a ratio of 2:1, keratinocytes to melanocytes and the cells were left in co-culture for 10 days, after which they were processed as described below. For immunofluorescence experiments, the cells were plated on glass coverslips coated with 1 mg/mL collagen (Sigma, C7661).

Animal care, use, and experimental procedures were conducted in accordance with recommendations of the European Community (86/609/EEC) and Union (2010/63/UE) and the French National Committee (87/848). Animal care and use were approved by the ethics committee of the Curie Institute in compliance with the institutional guidelines.

### Transfection and infection of melanocyte cell lines

All cells were transfected using the Amaxa Cell Line Nucleofector Kit L (Lonza, VCA-1005) according to the manufacturer’s recommendations. Briefly, cells were trypsinized and counted, with 7.5×10^5^ cells resuspended in the Amaxa transfection reagent. Four micrograms of each respective plasmid or 100 pmol of siRNA was added to the mixture, which was then transferred to a cuvette and incubated in the Amaxa nucleofector. Following nucleofection, the cells were added to a six-well dish containing pre-warmed growth media. The following day, the cells were washed and incubated in fresh growth media supplemented with TPA. Transfections were assayed 48 hours following nucleofection. Control GFP and β-catenin-GFP tagged plasmids have been previously described ^13^. The Dynlt3-GFP (pCMV3-mDynlt3-GFPSpark) and the control GFP vectors (pCMV3-GFP) were purchased from Sino Biological (MG5A2492-ACG and CV026, respectively).

9v cells expressing a control shRNA or an shRNA against Dynlt3 (targeting sequence: 5’-ATC AGA TGT TGT ATC CCA A-3’) were produced using viral particles and Dharmacon SMARTvector plasmids (VSC11715 and V3SM11241-233912870, respectively). Following infection, cells were selected using 1 μg/mL puromycin.

### siRNA sequences

siRNA targeting mouse *β-catenin* and *Dynlt3* were purchased from Dharmacon as a SMARTpool mix of 4 sequences. The sequences of the siRNA are as follows: for *β-catenin*: 5’-GAA CGC AGC AGC AGU UUG U-3’, 5’-CAG CUG GCC UGG UUU GAU A-3’, 5’-GCA AGU AGC UGA UAU UGA C-3’, 5’-GAU CUU AGC UUA UGG CAA U-3’; and for Dynlt3: 5’-CCC AUA AUA UAG UCA AAG A-3’, 5’-GGU GGU AAC GAU UAU AAU G-3’, 5’-GGG GAA AGC UUA CAA GUA C-3’, 5’-CAG AGG AGC CCG UAU GGA U-3’. A random, scrambled sequence (5’-AAU UCU CCG AAC GUG UCA CGU-3’) was used as a control.

### Semi-quantitative Real-time PCR

Total RNA was extracted from cells using the miRNeasy kit from Qiagen (217004) according to the manufacturer’s protocol. Reverse transcription reactions were performed with 500 ng of total RNA and M-MLV reverse transcriptase (Invitrogen), according to the manufacturer’s instructions. The cDNA generated was then subjected to quantitative real-time PCR for the analysis of gene expression using ABI 7900HT. Oligonulceotides used for PCR were as follows: for *β-catenin*, 5’-CGT GGA CAA TGG CTA CTC AA-3’ (F) and 5’-TGT CAG CTC AGG AAT TGC AC-3’ (R); for *Dynlt3* 5’-GCG ATG AGG TTG GCT TCA ATG CTG-3’ (F) and 5’-CAC TGC ACA GGT CAC AAT GTA CTT G-3’ (R); and for *Gapdh* 5’-ACC CAG AAG ACT GTG GAT GG-3’ (F) and 5’-CAC ATT GGG GGT AGG AAC AC-3’ (R). F means forward. R means reverse.

### Western blot analysis

Whole cell lysates were prepared from melanocyte cell lines using ice-cold RIPA buffer supplemented with complete protease inhibitor cocktail and PhosStop phosphatase inhibitor cocktail (Roche). For western blotting, 25 μg total protein was separated on 15% denaturing acrylamide SDS-PAGE gels and the proteins transferred to nitrocellulose membranes. Membranes were blocked in 5% non-fat milk in Tris-buffered saline supplemented with 0.05% Tween-20 (TBST) and probed with primary antibodies overnight. The signal was detected using peroxidase-conjugated anti-mouse or anti-rabbit secondary antibodies (refs) and enhanced chemiluminescence (ECL; Thermo Fisher). The primary antibodies were used as follows: Dynlt3, 1:200 (Sigma, hpa003938), β-catenin, 1:2000 (Abcam, ab6302), GFP, 1:500 (Institut Curie), and β-actin, 1:10,000 (Sigma, A5441).

### Immunofluorescence microscopy

Melanocyte cells were seeded on 18mm glass cover slips and grown to confluence, after which they were washed twice with ice-cold PBS and fixed-permeabilized with ice-cold methanol-acetone (1:1 solution) on ice for 5 minutes. Next, cells were washed twice with cold PBS and blocked with a blocking solution containing 1% BSA (w/v) and 10% FBS in PBS for 1 hour at room temperature. Cells were washed again twice with PBS and then incubated with primary antibodies against anti-Tyrp1 (Abcam, ab3312) and anti-GFP (Institut Curie) at 4°C overnight. The following day, the cells were washed again three times with PBS and then incubated with Alexa 555 anti-rabbit (ThermoFisher, A231572) or Alexa 488 anti-mouse (ThermoFisher, A21202) secondary antibodies for 1 hour at room temperature in the absence of light. Cells were then again washed two times before being counterstained with 0.5 µg/µL DAPI to visualize the nucleus. Coverslips were then mounted on glass slides using ProLong Gold antifade reagent (Invitrogen, P36934). Images were then taken using an Upright widefield Leica Microscope at 63X.

For co-culture experiments, mouse-keratinocyte co-cultures were washed twice with cold PBS and then fixed with 4% paraformaldehyde (VWR 1.04005.1000) for 20 min at room temperature. Following two washes with PBS, the cells were permeabilized with 0.2% v/v Triton X-100 in PBS for 10 min at room temperature. The cells were once again washed and then processed as described above, using anti-Hmb45 (Abcam, 787) and anti-plakoglobin (Abcam, 184919) primary antibodies. Co-cultures were examined using a CLSM SP5 Leica confocal microscope.

### Video microscopy and melanosome tracking

Cells were seeded in 6-well glass plates (MatTek Corporation P06G-1.5-20-F) and were imaged using the Ratio 2 inverted video microscope. Cells were maintained at 37°C in a humidified atmosphere of 5% CO_2_ while videos were being taken. Videos were taken at 100X objective over a period of 5 minutes, with one image taken every 0.5 sec, for a total number of 601 images. Videos were taken from a minimum of nine melanocytes per cell line/transfection from three independent experiments. From these melanocytes, the trajectories of 75 pigmented melanosomes were manually followed using the Manual Tracking plugin for *ImageJ* ^29^. Each melanosome path was chosen at random. The Manual Tracking allowed provided a number of different parameters that were used to characterize the movement of the individual melanosomes. Namely, we calculated the total distance the melanosomes moved, which was the sum of values for all of the 601 frames. The Euclidean distance was calculated as the distance between the melanosome position at the first frame and the last frame. The average distance was calculated as the total distance divided by the number of frames (601). Similarly, the average velocity was calculated by taking the average distance per frame and dividing by the time (0.5 sec). Finally, the percentage of time the melanosomes were paused was calculated by assessing the number of frames (out of the 601 total frames) where a “0” value was recorded for the distance travelled. Distribution of spatiotemporal descriptors of trajectories, in particular Mean Square Displacement characteristics, was computed and compared to simulated trajectories in order to distinguish convective *vs.* non-convective movements (^18^ and supplementary information).

### Melanosome number and localization quantification

Melanosome quantification was performed using ImageJ Analyze Particle feature. To exclude melanosomes nearby the nuclear region, a circle of 15μm was designated around the nucleus, as an arbitrary region of separation.

Melanosome localization in the different cell lines was quantified using ImageJ. A line was drawn along the length of each individual cell, with the width of the line extending to cover the entire cell. The pixel intensity along the length of the line was used to give a quantitative measure of melanosome distribution. Pixel intensity was measured from one end of the cell until reaching the nucleus, where the measurement stopped and was then continued on the other side of the nucleus. The data is presented as a measured pixel intensity over the percentage of the cell covered (as such, all measurements end at 100, regardless of the cell length). A trendline is presented (in red) to emphasize the trend.

### Flow Cytometry

For all flow cytometry experiments, cells were trypsinized and the resulting cell pellets were resuspended and fixed in 4% paraformaldehyde for 20 minutes at room temperature. Following centrifugation, the cell pellets were then washed twice with PBS containing 1% BSA and 0.1% saponin (Sigma, S4521). Cells were then incubated in primary antibodies against Hmb45 and plakoglobin (both at 1:100) for 2 hours at 4°C in the absence of light. Next, cell pellets were washed once with PBS/BSA/saponin and incubated in Alexa 647 mouse (ThermoFisher A31573) and Alexa 488 mouse secondary antibodies (at 1:500) for 1 hour. Cells were then washed in PBS and resuspended in PBS + 0.1% azide. Flow cytometric analyses were performed using a BD FACSCanto II machine.

### Melanosome acidity

Cells were seeded in 60 mm dishes and allowed to reach approximately 75% confluence. Acridine orange (Sigma, A6014) was added to the cells at 2.5 μg/mL and incubated for 20 minutes at 37°C. The cells were then incubated in warm PBS and imaged using a confocal microscope. To look at acridine orange staining, excitation wavelengths between 450-490 nm and emission between 590-650 nm were used ^30^. Melanosome acidity was assessed as by counting the number of pigmented melanosomes that were positive for acridine orange staining. This was done by merging the brightfield and acridine orange signals. The number of melanocytes assessed per field of view was always kept constant.

### Electron microscopy

Melanocytes seeded on coverslips were chemically fixed in 2.5 % (v/v) glutaraldehyde, 2 % (v/v) paraformaldehyde in 0.1M cacodylate buffer for 24h at 4°C, post-fixed with 1% (w/v) osmium tetroxide supplemented with 1.5% (w/v) potassium ferrocyanide, dehydrated in ethanol and embedded in Epon as previously described ^31^. Ultrathin sections of cell monolayers or tissue were prepared with a Reichert UltracutS ultramicrotome (Leica Microsystems) and contrasted with uranyl acetate and lead citrate. Electron micrographs were acquired on a Tecnai Spirit electron microscope (FEI, Eindhoven, The Netherlands) equipped with a 4k CCD camera (EMSIS GmbH, Münster, Germany).

## ACKNOWLEDGEMENTS

ZA was supported by the Fondation ARC and Labex ANR Labex CelTisPhyBio (ANR-11-LABX-0038 and ANR-10-IDEX-0001-02 PSL). ACP was supported by a fellowship from Institut Curie. We are grateful to the Rimbaud and Delmas families for their donations to our laboratory. This work was supported by Ligue Contre le Cancer, Fondation ARC, Institut Carnot, INCa, and ITMO Cancer, CNRS, INSERM and the French National Research Agency through the “Investments for the Future” program (France-BioImaging, ANR-10-INSB-04), and is under the program «Investissements d’Avenir» launched by the French Government and implemented by ANR Labex CelTisPhyBio (ANR-11-LABX-0038 and ANR-10-IDEX-0001-02 PSL).

We acknowledge the PICT-IBiSA, member of the France-BioImaging national research infrastructure.

## Supplementary information

This part is associated with Figure 5. Eight descriptors were defined: α, ρ, μ_2L_, μ_2S_, μ_3L_, μ_3S,_ μ_4L_ and μ_4S_. The comparison between a set of experiments and a set of simulated trajectories was performed by fitting the distributions of eight descriptors (MSD exponent, anisotropy, the second [q=2], third [q=3, skewness] and fourth [q=4, kurtosis] normalized moments projected on the two main L and S axis (Fig. 6 and Supplementary Fig. 5). The seven other descriptors refer only to 2D spatial characteristics and were obtained by projecting trajectory points on its own axis system, giving two unidimensional distributions which were statistically described by the computation of moments (E) of order q (2-4) (l corresponds to the center scalar distribution of the projection of the long axis, n to sample size, the second moment (when q=2) E2 [long axis] is named μ_2L-abs_), and by anisotropy defined as the ratio of the second moments μ_2S-abs_ / μ_2L-abs_ (s means short axis).

Small amplitude WT melanosome subpopulation was non convective (fitting with anisotropic Brownian movement + elastic tethering + noise), estimated to 39% of WT melanosome population. In contrast, only half of the population of the bcat* “small” was non-convective, and the other half was convective, estimated to 7% of bcat* population.

Large amplitude WT and bcat* subpopulations correspond to convective behavior of melanosomes. Convective behavior may correspond to two types of movement: intermittent or uniform. We consider the movement uniform when the velocity of each melanosome is constant during the whole observation period, and intermittent if the conditions are not fulfilled. We cannot discriminate between the uniform and intermittent behaviors of the wt or bcat* melanosomes. This is due to the fact the velocity is too low compared to the diffusion.

**Supplementary Table 1.** Mouse models associated with mutations in 67 genes involved in trafficking of melanosomes and other lysosome related organelles (LRO).

| Gene | Gene Product | Human Disease | Mouse Model | Stage |
| --- | --- | --- | --- | --- |
| **TYR** | TYR | OCA1 | Albino | Maturation |
| **OCA2** | OCA2/P | OCA2 | Pink eye dilution | Maturation |
| **TYRP1** | TYRP-1 | OCA3 | Brown | Maturation |
| **SLC45A2** | SLC45A2/MATP | OCA4 | Underwhite | Maturation |
| **DCT** | DCT/TRP-2 |  | Slaty | Maturation |
| **PMEL** | PMEL/Pmel17/Silver/gp100 |  | Silver | Maturation |
| **GPR143** | GPR143/OA1 | OA1 | Oa1 | Biogenesis |
| **MLANA** | MART1/melan-a |  | Melan-a | Maturation |
| **AP3B1** | AP-3β3a subunit | HPS2 | Pearl | Transport |
| **AP3D1** | AP-3 δ subunit |  | Mocha | Transport |
| **AP3M1** | AP-3 μ3a subunit |  | no<sup>2</sup> | Transport |
| **AP3S1** | AP-3 σ3a subunit |  | no<sup>2</sup> | Transport |
| **BLOC1S5** | Muted |  | Muted | Transport |
| **BLOC1S4** | Cappuccino | HPS1 | Cappuccino | Transport |
| **DTNBP1** | Dysbindin | HPS7 | Sandy | Maturation |
| **BLOC1S3** | BLOS3 | HPS8 | Reduced pigmentation | Biogenesis |
| **PLDN** | Pallidin | HPS9 | Pallid | Maturation |
| **BLOC1S1** | BLOS1 |  | No reported coat color | Transport |
| **BLOC1S2** | BLOS2 |  | no<sup>2</sup> | Transport |
| **SNAPIN** | Snapin |  | Perinatal lethality<sup>1</sup> | Transport |
| **HPS1** | HPS1 | HPS1 | Pale ear | Biogenesis |
| **HPS3** | HPS3 | HPS3 | Cocoa | Biogenesis |
| **HPS4** | HPS4 | HPS4 | Light ear | Biogenesis |
| **HPS5** | HPS5 | HPS5 | Ruby-eye 2 | Biogenesis |
| **HPS6** | HPS6 | HPS6 | Ruby-eye | Biogenesis |
| **VPS11** | VPS11 |  | no<sup>2</sup> | Transport |
| **VPS16** | VPS16 |  | no<sup>2</sup> | Transport |
| **VPS18** | VPS18 |  | no reported coat color<sup>3</sup> | Transport |
| **VPS33A** | VPS33A |  | Buff | Maturation |
| **VPS39** | VPS39/Vam6 |  | no<sup>2</sup> | Transport |
| **VPS41** | VPS41 |  | Embryonic lethal<sup>3</sup> | Transport |
| **VPS33B** | VPS33B | ARCS1 | no<sup>2</sup> | Maturation |
| **C14ORF13**<br>**3** | VIPAR/VPS16B | ARCS2 | no<sup>2</sup> | Transport |
| **RABGGTA** | Subunit of Rab geranylgeranyl transferase II |  | Gunmetal | Biogenesis |
| **LYST** | LYST | CHS | Beige | Transport |
| **MYO5A** | Myosin VA | GS1 | Dilute | Transport |
| **RAB27A** | RAB27A | GS2 | Ashen | Transport |
| **MLPH** | Melanophilin | GS3 | Leaden | Transport |
| **STX17** | Syntaxin 17 |  | no<sup>2</sup> | Elimination |
| **TRAPPC6A** | TRAPPC6A |  | Mhyp | Transport |
| **VAC14** | VAC14/ARPIKFYVE |  | Inglis (infantile gliosis) | Transport |
| **FIG4** | Fig/SAC3 | CMT4J | Pale tremor | Maturation |
| **AP1S1** | AP-1 σ1 subunit | | no<sup>2</sup> | Transport |
| **AP1S2** | AP-1 σ2 subunit | | no<sup>2</sup> | Transport |
| **AP1S3** | AP-1 σ3 subunit | | no<sup>2</sup> | Transport |
| **AP1M2** | AP-1 μ2 subunit | | no<sup>2</sup> | Transport |
| **ATG7** | Autophagy related 7 |  | Perinatal lethality<sup>3</sup> | Elimination |
| **ATG13** | Autophagy related 13 |  | no<sup>1</sup> | Elimination |
| **BACE2** | Bace2/ASP1 |  | No phenotype | Biogenesis |
| **CD63** | Cd63 |  | No coat color phenotype | Biogenesis |
| **DCTN1** | Dynactin 1 |  | Embryonic lethal<sup>3</sup> | Transport |
| **F2RL1** | F2RL1 |  | No coat color phenotype | Transfer |
| **GPNMB** | GPNMB |  | Eye color phenotype | Transfer |
| **KIF13A** | KIF13A |  | No coat color phenotype | Maturation/Transport |
| **KIF3A** | KIF3A |  | Embryonic lethal<sup>3</sup> | Transport |
| **KIF3B** | KIF3B |  | Embryonic lethal<sup>3</sup> | Transport |
| **KIF5B** | KIF5B |  | Embryonic lethal<sup>3</sup> | Transport |
| **MREG** | Melanoregulin |  | Dilute suppressor | Transfer |
| **RAB32** | RAB32 |  | no<sup>2</sup> | Biogenesis |
| **RAB11B** | RAB11B |  | no<sup>2</sup> | Transfer |
| **RAB38** | RAB38 |  | chocolate | Biogenesis |
| **RILP** | RILP |  | no<sup>2</sup> | Transport |
| **SLC24A5** | OCA6 |  | hypopigmentation | Maturation |
| **TSG101** | TSG101/VPS23 |  | Embryonic lethal<sup>3</sup> | Maturation |
| **MYO6** | Myosin VI |  | no<sup>2</sup> | Biogenesis |
| **DYNLT3** | DYNLT3 |  | no<sup>2</sup> | Transport |
| **RAB3D** | RAB3D |  | No coat color phenotype | Elimination |

| Gene | Gene Product | Human Disease | Mouse Model | Stage |
| --- | --- | --- | --- | --- |
| **TYR** | TYR | OCA1 | Albino | Maturation |
| **OCA2** | OCA2/P | OCA2 | Pink eye dilution | Maturation |
| **TYRP1** | TYRP-1 | OCA3 | Brown | Maturation |
| **SLC45A2** | SLC45A2/MATP | OCA4 | Underwhite | Maturation |
| **DCT** | DCT/TRP-2 |  | Slaty | Maturation |
| **PMEL** | PMEL/Pmel17/Silver/gp100 |  | Silver | Maturation |
| **GPR143** | GPR143/OA1 | OA1 | Oa1 | Biogenesis |
| **MLANA** | MART1/melan-a |  | Melan-a | Maturation |
| **AP3B1** | AP-3β3a subunit | HPS2 | Pearl | Transport |
| **AP3D1** | AP3 δ subunit |  | Mocha | Transport |
| **AP3M1** | AP-3 μ3a subunit |  | no<sup>2</sup> | Transport |
| **AP3S1** | AP-3 σ3a subunit |  | no<sup>2</sup> | Transport |
| **BLOC1S5** | Muted |  | Muted | Transport |
| **BLOC1S4** | Cappuccino | HPS1 | Cappuccino | Transport |
| **DTNBP1** | Dysbindin | HPS7 | Sandy | Maturation |
| **BLOC1S3** | BLOS3 | HPS8 | Reduced pigmentation | Biogenesis |
| **PLDN** | Pallidin | HPS9 | Pallid | Maturation |
| **BLOC1S1** | BLOS1 |  | No reported coat color | Transport |
| **BLOC1S2** | BLOS2 |  | no<sup>2</sup> | Transport |
| **SNAPIN** | Snapin |  | Perinatal lethality<sup>1</sup> | Transport |
| **HPS1** | HPS1 | HPS1 | Pale ear | Biogenesis |
| **HPS3** | HPS3 | HPS3 | Cocoa | Biogenesis |
| **HPS4** | HPS4 | HPS4 | Light ear | Biogenesis |
| **HPS5** | HPS5 | HPS5 | Ruby-eye 2 | Biogenesis |
| **HPS6** | HPS6 | HPS6 | Ruby-eye | Biogenesis |
| **VPS11** | VPS11 |  | no<sup>2</sup> | Transport |
| **VPS16** | VPS16 |  | no<sup>2</sup> | Transport |
| **VPS18** | VPS18 |  | no reported coat color<sup>3</sup> | Transport |
| **VPS33A** | VPS33A |  | Buff | Maturation |
| **VPS39** | VPS39/Vam6 |  | no<sup>2</sup> | Transport |
| **VPS41** | VPS41 |  | Embryonic lethal<sup>3</sup> | Transport |
| **VPS33B** | VPS33B | ARCS1 | no<sup>2</sup> | Maturation |
| **C14ORF13**<br>3 | VIPAR/VPS16B | ARCS2 | no<sup>2</sup> | Transport |
| **RABGGTA** | Subunit of Rab geranylgeranyl transferase II |  | Gunmetal | Biogenesis |
| **LYST** | LYST | CHS | Beige | Transport |
| **MYO5A** | Myosin VA | GS1 | Dilute | Transport |
| **RAB27A** | RAB27A | GS2 | Ashen | Transport |

**Supplementary Figure 1.**
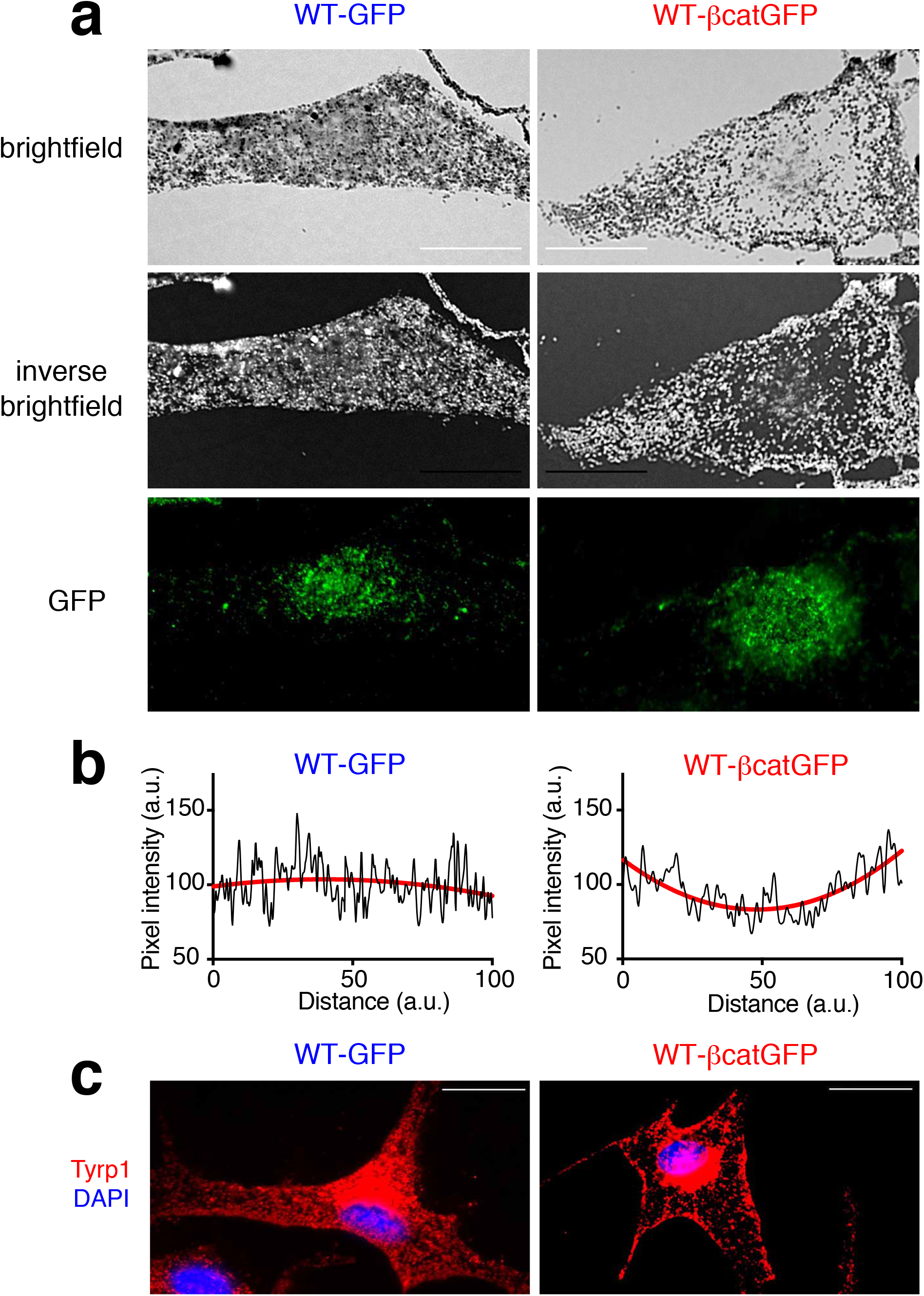
Melanosomes are distributed at the periphery of melanocytes expressing exogenous activated β-catenin. a) Brightfield (top), inverse brightfield (middle) and GFP (bottom) images of WT (9v) melanocytes transiently expressing GFP or GFP-tagged β-catenin. Bar, 20 µm. b) Quantitation of pixel intensity (in black) across the length of the WT-GFP and WT-β-catGFP melanocytes shown in Panel A. Trendlines are shown in red. c) Immunofluorescence analysis of WT melanocytes transiently expressing GFP or GFP-tagged b-catenin. Cells were processed for immunofluorescence and stained with anti-Tyrp1 (red) antibodies and the nuclei were stained with DAPI (blue). Bar, 20 µm.

**Supplementary Figure 2.**
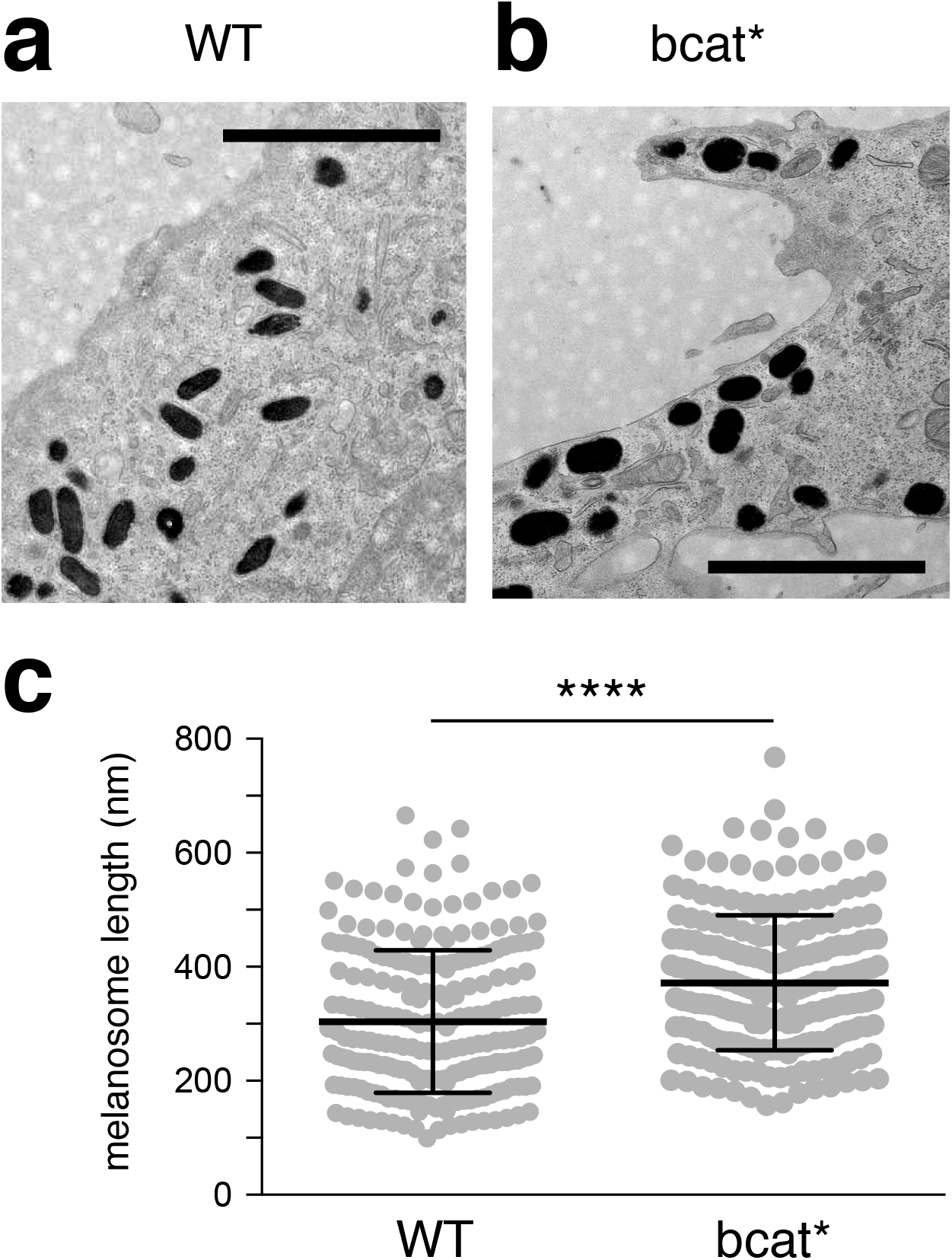
bcat* melanosomes are larger than WT melanosomes. a,b) Transmission electron microscopy of WT (a) and bcat* (b) melanocytes. Scale bar: 2 μm. c) Length of melanosome was determined from four WT and five bcat* melanocytes with a total of at least 200 melanosomes per cell line. Statistical significance was measured using two-sided Mann-Whitney test. **** p < 0.0001.

**Supplementary Figure 3.**
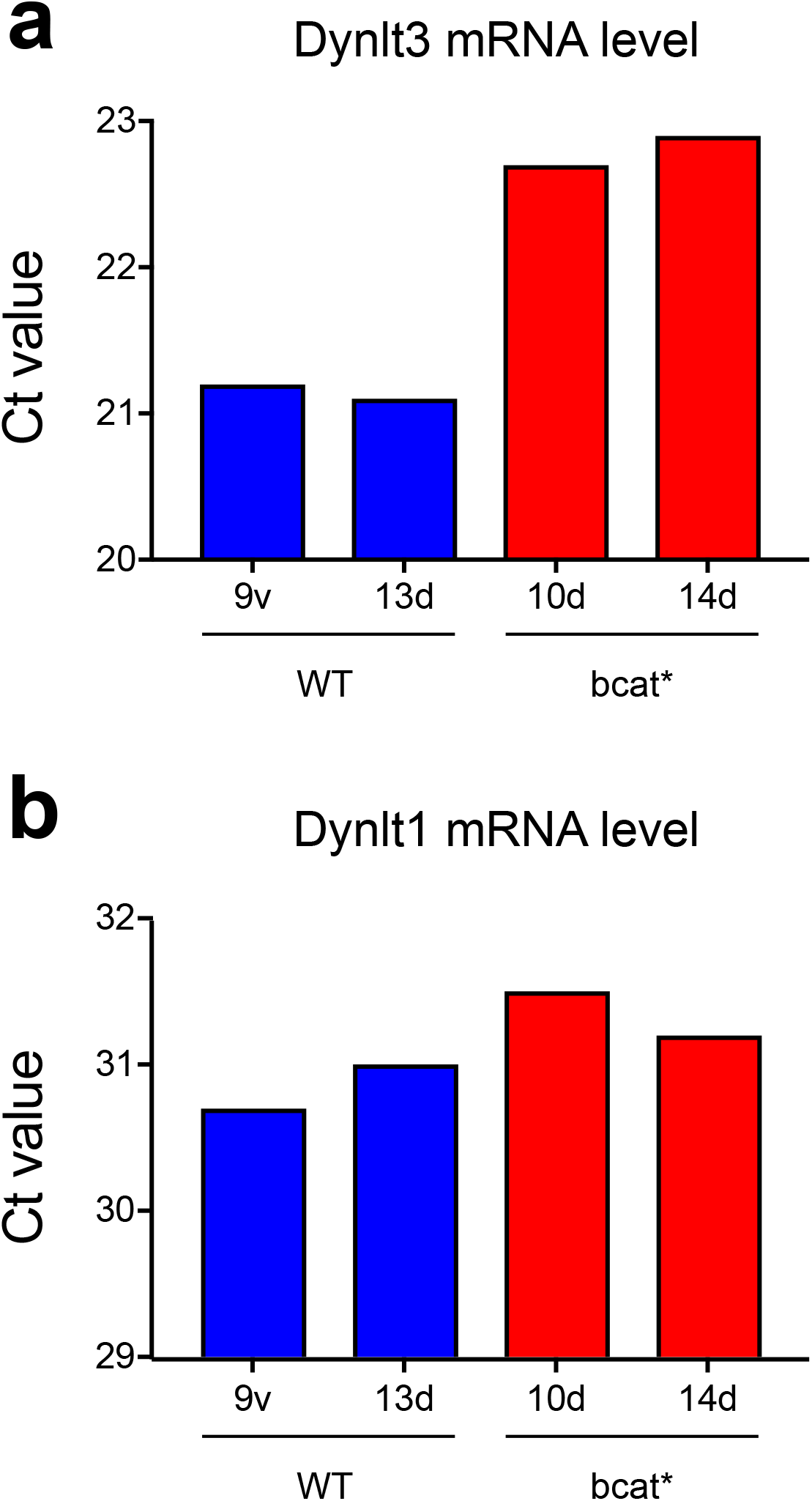
Dynlt3, and not Dynlt1, is expressed in WT and downregulated in bcat* melanocytes. Ct values for RT-qPCR analysis of Dynlt3 (A) and Dynlt1 (B) mRNA expression in two cell lines derived from WT mice (9v and 13d) and two from bcat* mice (10d and 14d).

**Supplementary Figure 4.**
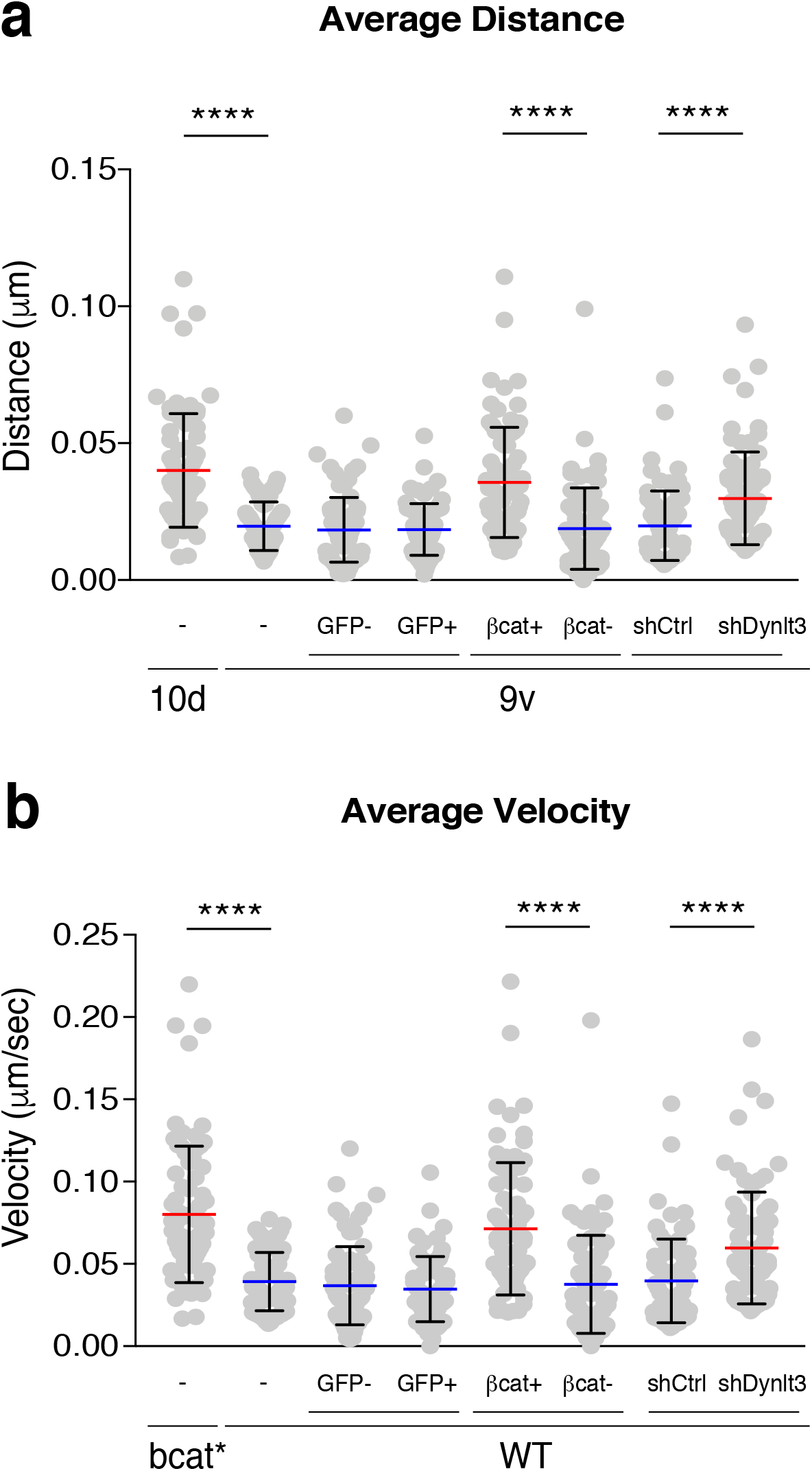
Melanosome movement is increased in cells producing active β-catenin or with diminished level of Dynlt3. Melanosome movement (corresponding mainly to stage IV melanosome) was assessed by brightfield video microscopy over a period of 5 minutes and melanosome trajectories were followed using *ImageJ* software. For each cell line, 75 melanosomes were followed from a minimum of five independent cells. WT (9v) cell lines were transfected with control GFP or β-catenin-GFP expression vectors, respectively. Analyses were performed on both transfected (i.e. GFP-positive) and non-transfected (i.e. GFP-negative) cells. In addition, WT cells were transfected with either control or Dynlt3 shRNA vectors and analyses were performed on the resulting red cells. Each dot represents one melanosome. The average distance (a) and the average velocity (b) for each melanosome are shown, Statistical significance was measured using two-sided Mann-Whitney test. **** p < 0.0001.

**Supplementary Figure 5.**
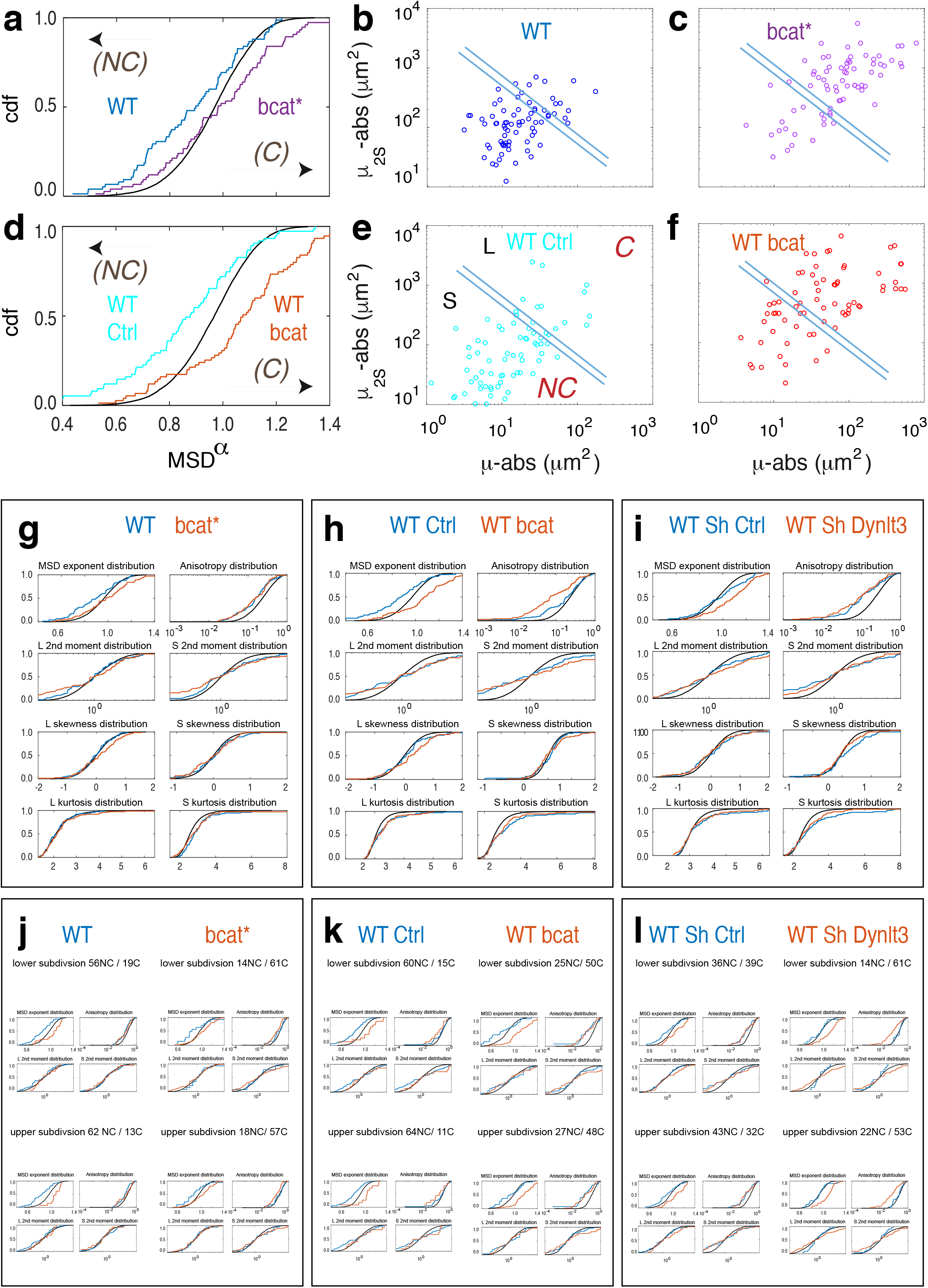
Mathematical modeling of melanosome trajectories. a,d) α MSD distributions for 9v and 10d (a), and 9v-Ctrl and 9v-bcat (d) populations of trajectories, compared to the distribution for simulated Brownian (purely diffusive – black line) random walks (nwalks=10 000, with the same nsteps (601) as for experimental trajectories). Theoretical shifts compared to Brownian process when adding convection, noise, tethering or confinement are indicated with arrows. b,c,e,f) µ_2S_, µ_2L_ plots reflecting the extent of 9v (b), 10d (c), 9v Ctrl (e), and 9v bcat (f) trajectories. Each point corresponds to one trajectory. Lines correspond to frontiers between non-convective (NC) and convective (C) trajectories: see SI for criteria and validation. g,h,i) α MSD, anisotropy, L and S second moments, L and S skewness, and L and S kurtosis distributions for 9v and 10d (g), 9v-Ctrl and 9v-bcat (h), and 9v Sh Ctrl and 9v Sh Dynlt3 (i) populations of trajectories, compared to the distribution for simulated Brownian (purely diffusive – black line). j,k,l) Criteria to establish the frontiers for non-convective (NC) and convective (C) movement for α MSD, anisotropy, and L and S second moments distributions. On the top and the bottom the lower and upper subdivisions are given, respectively.

**Supplementary Figure 6.**
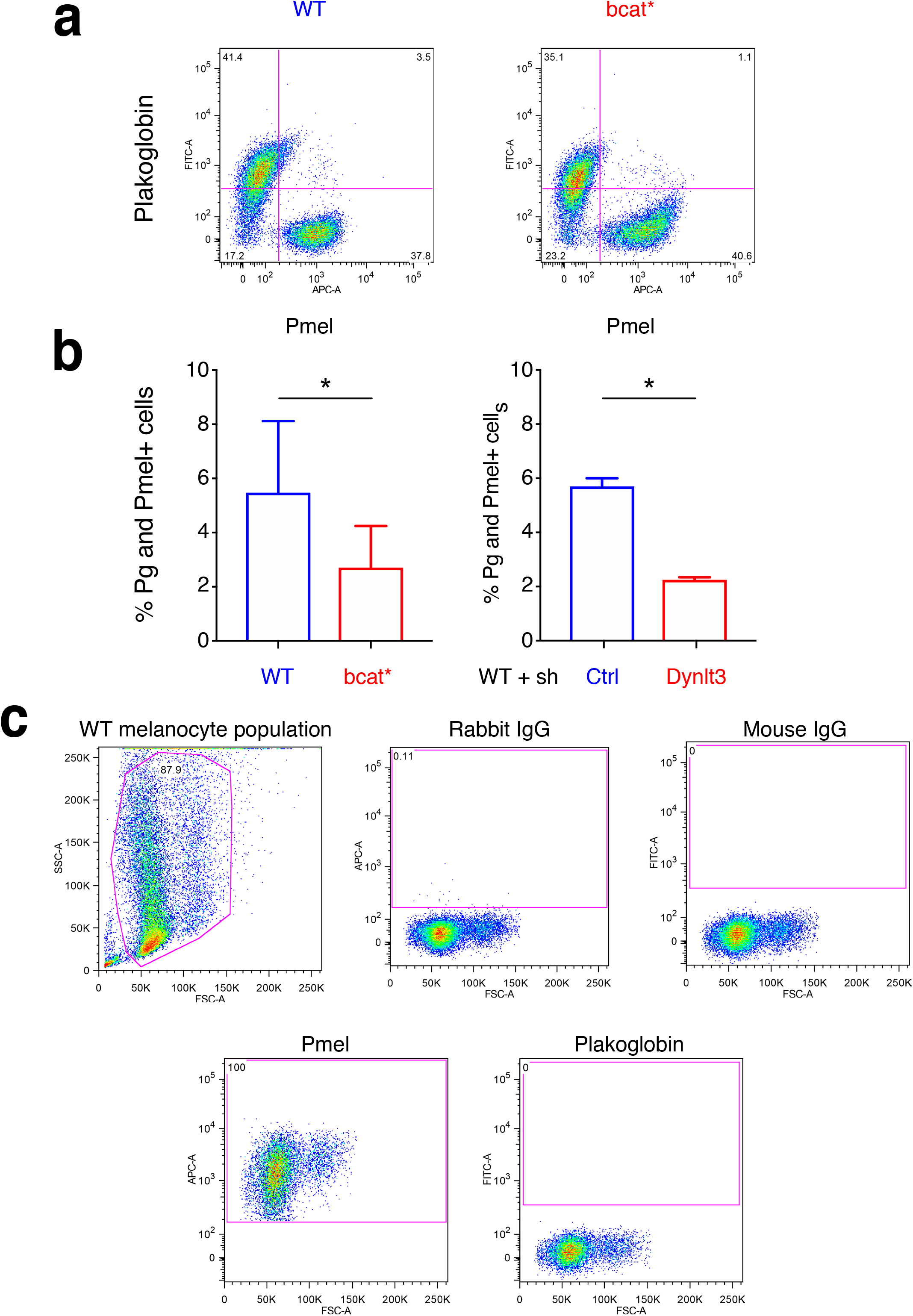
FACS analysis of melanosome transfer. a) FACS analysis of Hmb45 and plakoglobin positive cell populations in WT and bcat* melanocytes. b) Quantitation of melanosome transfer assayed by FACS. WT (9v), bcat* (10d), WT + sh Ctrl or WT + sh Dynlt3 melanocytes were co-cultured with Balb/c MK mouse keratinocytes for 10 days, after which the co-cultures were fixed and processed for FACS analyses using Hmb45 and Plakoglobin antibodies. The percentage of Plakoglobin and Hmb45 double positive cells within each respective co-culture is presented. Quantitation was done from a minimum of five independent co-culture experiments. c) Gating strategy for FACS experiments. Melanocyte and keratinocyte cells were stained with either control IgG, plakoglobin or Pmel antibodies. The live cell population was chosen for gating. IgG control antibodies were used to establish a baseline of background signal. Following this, the established gate was applied to samples stained with the respective antibodies of interest. The gates were kept constant for all cell lines (melanocyte and keratinocyte) and for the co-culture experiments.

